# PhysiCell: an Open Source Physics-Based Cell Simulator for 3-D Multicellular Systems

**DOI:** 10.1101/088773

**Authors:** Ahmadreza Ghaffarizadeh, Randy Heiland, Samuel H. Friedman, Shannon M Mumenthaler, Paul Macklin

## Abstract

Many multicellular systems problems can only be understood by studying how cells move, grow, divide, interact, and die. Tissue-scale dynamics emerge from systems of many interacting cells as they respond to and influence their microenvironment. The ideal “virtual laboratory” for such multicellular systems simulates both the biochemical microenvironment (the “stage”) and many mechanically and biochemically interacting cells (the “players” upon the stage).

PhysiCell—physics-based multicellular simulator—is an open source agent-based simulator that provides both the stage and the players for studying many interacting cells in dynamic tissue microenvironments. It builds upon a multi-substrate biotransport solver to link cell phenotype to multiple diffusing substrates and signaling factors. It includes biologically-driven sub-models for cell cycling, apoptosis, necrosis, solid and fluid volume changes, mechanics, and motility “out of the box.” The C++ code has minimal dependencies, making it simple to maintain and deploy across platforms. PhysiCell has been parallelized with OpenMP, and its performance scales linearly with the number of cells. Simulations up to 10^5^-10^6^ cells are feasible on quad-core desktop workstations; larger simulations are attainable on single HPC compute nodes.

We demonstrate PhysiCell by simulating the impact of necrotic core biomechanics, 3-D geometry, and stochasticity on the dynamics of hanging drop tumor spheroids and ductal carcinoma in situ (DCIS) of the breast. We demonstrate stochastic motility, chemical and contact-based interaction of multiple cell types, and the extensibility of PhysiCell with examples in synthetic multicellular systems (a “cellular cargo delivery” system, with application to anti-cancer treatments), cancer heterogeneity, and cancer immunology. PhysiCell is a powerful multicellular systems simulator that will be continually improved with new capabilities and performance improvements. It also represents a significant independent code base for replicating results from other simulation platforms. The PhysiCell source code, examples, documentation, and support are available under the BSD license at http://PhysiCell.MathCancer.org and http://PhysiCell.sf.net.

**Author Summary:** This paper introduces PhysiCell: an open source, agent-based modeling framework for 3-D multicellular simulations. It includes a standard library of sub-models for cell fluid and solid volume changes, cycle progression, apoptosis, necrosis, mechanics, and motility. PhysiCell is directly coupled to a biotransport solver to simulate many diffusing substrates and cell-secreted signals. Each cell can dynamically update its phenotype based on its microenvironmental conditions. Users can customize or replace the included sub-models.

PhysiCell runs on a variety of platforms (Linux, OSX, and Windows) with few software dependencies. Its computational cost scales linearly in the number of cells. It is feasible to simulate 500,000 cells on quad-core desktop workstations, and millions of cells on single HPC compute nodes. We demonstrate PhysiCell by simulating the impact of necrotic core biomechanics, 3-D geometry, and stochasticity on hanging drop tumor spheroids (HDS) and ductal carcinoma in situ (DCIS) of the breast. We demonstrate contact- and chemokine-based interactions among multiple cell types with examples in synthetic multicellular bioengineering, cancer heterogeneity, and cancer immunology.

We developed PhysiCell to help the scientific community tackle multicellular systems biology problems involving many interacting cells in multi-substrate microenvironments. PhysiCell is also an independent, cross-platform codebase for replicating results from other simulators.

## Introduction

Many significant multicellular systems problems—such as tissue engineering, evolution in bacterial colonies, and tumor metastasis—can only be understood by studying how individual cells grow, divide, die, move, and interact [1–5]. Tissue-scale dynamics emerge as cells are influenced by biochemical and biophysical signals in the microenvironment, even as the cells continually remodel the microenvironment. Thus, the ideal “virtual laboratory” for multicellular systems biology must simultaneously simulate (1) tissue microenvironments with multiple diffusing chemical signals (e.g., oxygen, drugs, and signaling factors), and (2) the dynamics of many mechanically and biochemically interacting cells [5]. We recently published and open sourced the first part of such a platform: BioFVM, a biotransport solver that can efficiently simulate secretion, diffusion, uptake, and decay of multiple substrates in large 3-D microenvironments, even on desktop workstations [6]. We now introduce and release as open source PhysiCell: a mechanistic off-lattice agent-based model that extends BioFVM to simulate the tissue-scale behaviors that emerge from basic biological and biophysical cell processes.

### Prior work

Several major computational frameworks are available for studying 3-D multicellular systems. CompuCell3D [7] and Morpheus [8] use Cellular Potts methods to simulate cells and their morphologies. They are very user-friendly packages with graphical model editors, integrated ODE and PDE solvers, and support for molecular-scale sub-models, but they currently cannot scale to large numbers (10^5^ or more) of cells. TiSim (part of the CellSys package [9]) can simulate many more cells by using a cell-centered, off-lattice approach. However, it is currently closed source, and its executables are restricted to a limited set of simulation types. Chaste [10] is a powerful, well-developed framework for multicellular modeling with integrated PDE and ODE solvers, and both cell- and vertex-based simulations of 10^5^ or more cells. However, its complex codebase has many dependencies that can impede participation by new developers; it is only cross-platform compatible by virtual machines. Biocellion [11] can simulate billions of cells on cluster computers, but it is closed source, and its restrictive user license has hindered adoption. Timothy was recently developed to simulate large-scale colonies of 10^9^ cells on high-performance supercomputers [12,13]. Most of these platforms offer a general-purpose pre-compiled “client” that can load models and settings from an XML file; this helps overcome difficulties stemming from complex dependencies. See S1 Text for a detailed software comparison.

These platforms typically require users to write their own code for all cell activities, by scripting basic built-in functions. (e.g., build a cell cycle model from API functions to overwrite cell volume and duplicate agents when appropriate.) As configured “out of the box,” none have built-in models for cell cycling, apoptosis, and necrosis, even though these fundamental behaviors are needed in many multicellular simulations. Only CompuCell3D and Morpheus have built-in volume regulation features. Most of these packages include PDE solvers that can simulate 3-D biotransport of one or more diffusible factors in the microenvironment, but generally they are applied sequentially to one PDE at a time, meaning that solving 10 PDEs requires 10 times more computational work than solving a single PDE. This approach is not expected to efficiently scale to 3-D simulations with many diffusible factors—a key requirement in reconciling secretomics with single-cell and multicellular systems biology, particularly as we work to understand cell-cell communication involving many cell-secreted factors.

### Scientific design goals and use cases

We aim to create a framework for building multicellular simulations that investigate the relationship between (diffusional) substrate limitations, multicellular biochemical communication, and essential phenotypic processes: cell cycling/division, death, volume changes, motility, adhesion, and volume exclusion/”repulsion” [1]. Building upon our experiences in agent-based modeling [14–16], we seek a system with a minimal set of mechanics to avoid cell lattice effects [2]; hence we use an off-lattice model with basic cell adhesion and “repulsion” implemented as simple potential functions [9, 10,14]. Following our work in [14–16], we include basic models of gradual cell volume changes, rather than static cell volumes. This avoids non-physical “jumps” in cell velocity following cell division events: the sudden localized doubling of cell density causes cells to overlap, leading to large, temporary, and non-physical “repulsive” forces that can manifest as non-physical “tears” in simulated tissues.

While PhysiCell was originally developed for problems in cancer [14], its diffusional and phenotypic processes are not specific to cancer. Users can introduce new environmental substrates (e.g., extracellular matrix (ECM) as a substrate with zero diffusion coefficient), new cell types (e.g., fibroblasts with high motility, low proliferation, and secretion and degradation of ECM), or new systems of cells (e.g., a network of vascular agents that release oxygen as in [6], and that can divide and move along gradients of angiogenic growth factors). Modelers can also use PhysiCell’s core functions to create new libraries that simulate physiological systems, similar to how Microvessel Chaste [17] built upon the core functions of Chaste [10] to simulate new classes of problems in vascular remodeling. In Additional PhysiCell examples, we use PhysiCell’s basic building blocks to simulate synthetic multicellular systems, heterogeneous tumors, and the innate immune system as it interacts with tumor cells. As we continue to apply PhysiCell to our own work in tissue engineering, angiogenesis, microbial dynamics, and cancer, we plan to release new optional modules that introduce new capabilities and simplify modeling of *in vivo* tissues and other systems. See Availability and Future Directions.

Thus, PhysiCell is designed to be a general-purpose toolkit for exploring multicellular systems, and the multicellular behaviors that emerge from a user-implemented set of cell phenotypic rules (i.e., a set of hypotheses). Other potential uses include simulation-generated synthetic datasets that investigate the meaning and shortcomings of experimental measurements [18,19]. PhysiCell’s design has allowed straightforward deployment on supercomputers for high-throughput simulation studies. We recently explored over 250 instances of a 3-D cancer-immune model (see Additional PhysiCell examples) to study the impact of immune cell motility and cancer-immune adhesion dynamics on a cancer immune response, thus reducing 1.5 years’ worth of simulations to 2 days on a Cray supercomputer [20, 21]. We anticipate that users can similarly use PhysiCell to efficiently explore large spaces of parameter values and hypotheses (model rules) with biophysically realistic, 3-D models. In Fall 2017, we started using PhysiCell in undergraduate and graduate education, as a self-contained tool for computational tissue engineering and cancer systems engineering. Lastly, PhysiCell also serves as an independent codebase to cross-validate model predictions in Chaste, Biocellion, TiSim, Timothy, and other platforms.

### Computational design goals

PhysiCell aims to balance computational speed, built-in standard functionality, flexibility, and codebase simplicity. It includes a built-in library of standardized cell cycle and cell death models co-developed with biologists and modelers here and in the MultiCellDS standardization process [22,23], force-based cell-cell interaction mechanics, motility, and volume regulation. Users can replace any of these built-in models with their own, and they can dynamically assign custom functions to any agent at any time. Through BioFVM, PhysiCell can couple cell phenotype to many diffusible substrates. It is the only simulation package to explicitly model the cell’s fluid content—a key aspect in problems such as cryobiology [24]. It can simulate systems of 10^5^-10^6^ cells on desktop workstations, and 10^6^ or more cells on single HPC compute nodes. All this functionality and performance is achieved with only two external dependencies, and a fully crossplatform C++ codebase that we have compiled and tested on Linux, OSX, and Windows. We also distribute PhysiCell as a virtual appliance—with a full Linux desktop, 64-bit g++ compiler, and visualization tools—that can run in widespread virtual machine software like VirtualBox.

## Design and Implementation

PhysiCell is designed to study the dynamics and interactions of thousands or millions of cells in 3-D microenvironments, with microenvironment-dependent phenotypes. It uses a lattice-free, physics-based approach to reduce grid-based artifacts. It provides optimized, biologically realistic functions for key cell behaviors, including: cell cycling (multiple models for *in vitro* and *in vivo*-focused simulations) cell death (apoptosis and necrosis), volume regulation (fluid and solid biomass; nuclear and cytoplasmic sub-volumes), motility, and cell-cell mechanical interactions. Each cell agent is built with a hierarchical Phenotype data structure; key phenotypic processes are triggered and controlled by modifying the phenotype data. This allows users to focus on modeling microenvironment-dependent triggers of standard cell processes, rather than coding these basic processes. However, to maintain flexibility, PhysiCell is written in a modular manner so that users can extend, rewrite, or replace its functions. Users can also create custom rules, and assign them to individual agents. It is fully coupled to a fast multi-substrate diffusion code (BioFVM) that solves for vectors of diffusing substrates, so that users can tie cell phenotype to many diffusing signals.

We note that for many problems in cancer biology and tissue engineering that drove development of PhysiCell, diffusive biotransport occurs at relatively fast time scales (on the order of 0.1 min or faster) compared to cell mechanics (~ 1 min) and cell processes (~ 10 to 100 min or slower). PhysiCell takes advantage of this by using three separate time step sizes (Δ*t*_diff_, Δ*t*_mech_, and Δ*t*_cells_). In particular, the cell phenotypes and arrangement (operating on slow time scales) can be treated as quasi-static when advancing the solution to the biotransport PDEs, so BioFVM can be called without modification with the cell arrangements fixed. See Time steps for further discussion. We provide default time step sizes that should suffice for typical applications in cancer biology and tissue engineering. Problems with inherently different time scales (e.g., advection-dominated problems, or with fast-moving bacteria) will need to adjust the time steps accordingly to avoid spurious oscillations that would be expected if cell-based sources and sinks are moving relatively quickly.

PhysiCell was built by extending the `Basic_Agent` class in BioFVM [6] (a static, nonmoving object that can secrete and uptake substrates) into a fully dynamic Cell class with changing cell volume, cycle progression, death processes, motility, and mechanics. This allows the cells to directly and efficiently interface with the multi-substrate microenvironment. PhysiCell is written in cross-platform compatible C++ and is self-contained (with minimal dependencies). It can be compiled in any C++11 compiler with OpenMP support. This simplifies installation and improves the reproducibility of the experiments. We have tested PhysiCell on Windows through MinGW-w64, and on OSX and Linux via `g++`. (OSX users should note that Xcode does not include a compliant OpenMP-compatible `g++`; they should follow our tutorials for installing a compliant g++ via Homebrew or MacPorts. See http://MathCancer.org/blog/physicell-tutorials/.) PhysiCell’s only external dependencies are pugixml [25] (for XML parsing) and BioFVM [6] for 3-D multi-substrate diffusion. For the user’s convenience, compatible versions of pugixml and BioFVM are included in every download.

The code has been parallelized in OpenMP to make use of multi-core desktop workstations and single HPC compute nodes. In testing, its performance (the computational time to simulate a fixed amount of time on a fixed domain) scales linearly in the number of cells present in the simulation. Simulations of up to 10^6^ cells are feasible on desktop workstations, and simulations beyond 10^6^ cells are possible on typical HPC compute nodes.

### Overall program flow

After initializing the microenvironment (through BioFVM [6]) and cells, and initializing the current simulation time *t* = 0, PhysiCell tracks (internally) *t*_mech_ (the next time at which cell mechanics functions are run), *t*_cells_ (the next time at which cell processes are run), and *t*_save_ (the next simulation data output time), with output frequency Δ*t*_save_. Initially, we set:

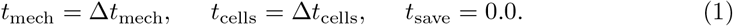

PhysiCell repeats the main program loop until reaching the maximum simulation time:

1. Save the simulation state if *t* ≥ *t*_save_. Set *t*_save_ = *t*_save_ + Δ*t*_save_.
2. Run BioFVM to update the biochemical microenvironment for cell-based secretions and uptake, and reaction-diffusion, for the current fixed cell positions. See Biochemical microenvironment.
3. For the fixed cell positions and chemical substrate fields, if *t* ≥ *t*_cells_, run the cell processes for each cell:
  (a) Update cell parameters using the cell’s `update_phenotype functions`. See Cell characteristics and state.
  (b) Advance the cell cycle (or death) model. See Cell cycling and Cell death
  (c) Advance the cell’s volume (and sub-volumes) using the `volume_update_function`. See Cell volume.
  Then set *t*_cells_ = *t*_cells_ + Δ*t*_cells_.
4. If *t* > *t*_mech_, then:

(a) For each cell, calculate the force-based cell velocities via `update_velocity`, add the contribution of motility (which runs `update_migration_bias` as needed), and run the custom function `custom_celljrule`. See Cell motility and Cell mechanics and motion.
(b) For each cell, update the position using the Adams-Bashforth method. See Cell mechanics and motion. Set *t*_mech_ = *t*_mech_ + Δ*t*_mech_.
5. Update the current simulation time by t = t + Δt_diff_. Return to Step 1.

Steps 3a-c can be combined in a parallelized OpenMP loop; we flag cells for division and removal and process these queues serially after the parallel loop to avoid data corruption. Step 4a can be parallelized across the cells by OpenMP (because cell velocities are location-dependent, and the cell positions are fixed throughout 4a), and then Step 4b can be parallelized across the cells with these computed velocities.

### Biochemical microenvironment

We use BioFVM to simulate the chemical microenvironment with a vector of reaction-diffusion PDEs with both bulk source/sinks and cell-centered sources and sinks [6]. To briefly summarize that prior work, we model the biochemical microenvironment (with computational domain Ω and boundary ∂Ω, discretized as a Cartesian mesh for computational efficiency) as a vector of reaction-diffusion PDEs for a vector of chemical substrates *ρ* of the form

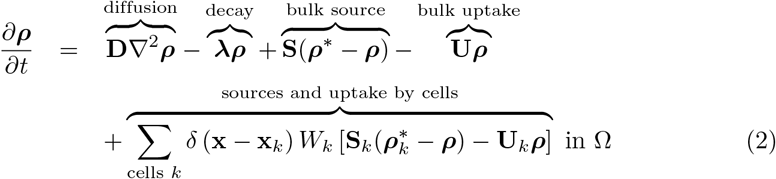

with zero flux conditions on ∂Ω. Here, *δ*(x) is the Dirac delta function, x_*k*_ is the *k*^th^ cell’s position, *W_k_* is its volume, **S**_*k*_ is its vector of source rates, **U**_*k*_ is its vector of uptake rates, and ***ρ***\* is the vector of saturation densities (the densities at which the cells stop secreting). Likewise, **D** and **λ** are the vectors of diffusion coefficients and decay rates, **S** is the bulk supply rate, and **U** is the bulk uptake function. All vector-vector products ab are element-wise (Hadamard product).

Numerically, we solve for the solution at time *t* + Δ*t*_diff_ by a first-order operator splitting: we first solve the bulk source/sink terms across the domain (and overwrite the stored solution), then solve the cell-centered source/sink terms (and overwrite the solution), and then the reaction-diffusion terms (again, overwriting the stored solutions). As we detailed and verified in [6], because we used (numerically stable) first-order backward differences in all our time discretizations, the overall method is first-order accurate in time and numerically stable.

After this operator splitting, the bulk source/sink terms are a decoupled set of systems of ODEs (one vector of ODEs in each computational voxel), which we solve by the backwards Euler method for first-order accuracy and numerical stability. This is trivially parallelized by OpenMP by dividing the voxels across the processor cores. Similarly, we solve the cell-based source/sink equations (one system of ODEs per cell) by backwards-Euler, overwriting the solution in the voxel containing the cell’s center. As we showed in [6], this is first-order accurate and numerically stable, and it performs best when the computational voxels are comparable to the cells’ sizes or larger. In all our work, we satisfy this requirement by using 20 *μ*m voxels. As with the bulk source/sinks, we trivially parallelize by dividing the cell source/sinks (one system of ODEs per cell) across the processor cores with OpenMP.

To solve the vector of diffusion-decay PDEs, we use the locally one-dimensional method: a specialized operator splitting that turns the 3-D PDEs into a sequence of one-dimensional PDEs. In each *x*-strip, *y*-strip, or *z*-strip, we discretize the PDEs with second-order centered differences (spatial derivatives) and first-order backwards differences (time derivatives) to obtain a tridiagonal linear system in each strip, which we solve exactly with a tailored Thomas solver. The Thomas solver itself cannot be trivially parallelized, but because we have many simultaneous *x*-, *y*-, and *z*-strips, we can distribute many instances of the Thomas solver across processor cores by OpenMP. This technique, along with other optimizations (tailored overloaded vector operators, a vectorized Thomas solver, pre-computing and caching the forward sweeps of the Thomas solvers, and other similar optimizations) was tested to scale linearly in the number of substrates (number of PDEs), the number of computational voxels, and the number of discrete cell sources/sinks. The method was numerically stable even for large Δ*t*, first-order accurate in time, and second-order accurate in space. Simulating 10 PDEs on 1,000,000 voxels takes approximately 2.6 times more computational time than simulating a single PDE. We found that for typical magnitudes of D, S, U, and λ, using Δ*t*_diff_ = 0.01 min and Δ*x* = 20 *μ*m gave solutions with 5% relative accuracy or better. For more algorithmic detail and extensive convergence testing on a variety of problems, see [6] and its supplementary material.

### Agent-based cell model

PhysiCell implements key cell-scale processes—cell cycling and death, volume changes, mechanics, and motility—and lets users link the parameters of these processes to microenvironmental conditions. PhysiCell then scales these basic cell-scale hypotheses to simulate thousands or millions of cells, from which tissue-scale behavior emerges. Here, we summarize the key functions. For each sub-model, see S1 Text for the full equations, further biological motivation, and reference parameter values.

#### Cell characteristics and state

Cell agents have a variety of phenotypic properties, including position (x_*i*_), volume (and sub-volumes), cell cycle or death status, and mechanics (adhesive, deformation, and motility) parameters. Each cell has a hierarchically-structured, independent `Phenotype` object that organizes information on cell cycling, death, volume, geometry, mechanics, motility, and secretion. Each cell also has an independent `Cell_Functions` object that collects functions to update cell volume, motility, velocity (mechanics), and other properties. See the User Guide (S2 Text, included in every PhysiCell download Version 1.2.0 or later) for a list of all the cell agents’ attributes and functions to access/update them. Below, we describe standardized, built-in models to update these properties. The models can be replaced by user-defined functions; the supplied models serve as biophysically reasonable default functions that capture the key aspects of these processes.

#### Cell volume

Each cell tracks *V* (total volume), *V*_F_ (total fluid volume), *V*_S_ (total solid volume), *V*_NS_ (nuclear solid volume), *V*_CS_ (cytoplasmic solid volume), *V*_N_ (total nuclear volume), and *V*_C_ (total cytoplasmic volume). Key parameters include nuclear solid, cytoplasmic solid, and fluid rate change parameters (*r*_N_, *r*_C_, and *r*_F_), the cell’s “target” fluid fraction *f*_F_, target solid volume 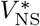, and target cytoplasmic to nuclear volume ratio *f*_CN_. For each cell, these volumes are modeled with a system of ODEs:

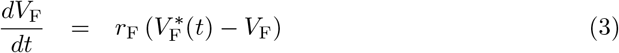

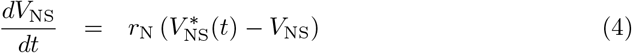

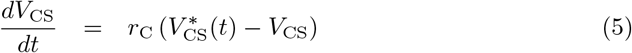

where we use the constitutive laws

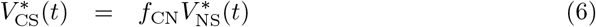

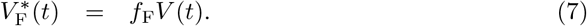

These parameters are updated as the cell progresses through its current cycle or death process. (See Cell cycling and Cell death.) For example, we halve 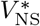 after a cell division and double it upon re-entry to the cell cycle. We set 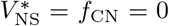 at the onset of apoptosis, and we set *r*_F_, *r*_C_, and *r*_N_ based on key apoptosis time scales. See S1 Text for the full equations, sample solution plots, further biological motivation, convergence testing, and reference parameter values. These ODEs are numerically solved using first-order forward Euler discretizations.

#### Cell motility

Each cell can set its own persistence time (*T*_per_), migration speed (*s*_mot_), migration bias direction (d_bias_, a unit vector), and a migration bias *b* that ranges from 0 (Brownian motion) to 1 (deterministic motion along d_bias_). When updating the cell’s velocity, its migration velocity *V*_mot_ is added to the currently velocity (as calculated by mechanics; see Cell mechanics and motion). The cell changes its migration velocity stochastically between *t* and *t* + Δ*t*_mech_ with probability

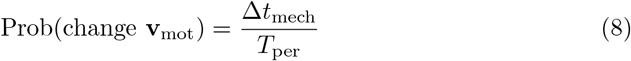

The user-defined function `update_migration_bias` (see the User Guide S2 Text and S1 Text) sets v_bias_, *b*, and *s*_mot_. The migration velocity v_mot_ is then updated according to

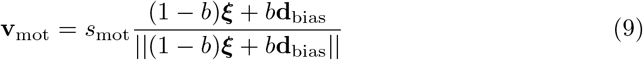

where *ξ* is a random unit vector. See the user manual (S2 Text) and S1 Text for more details, as well as examples for biased migration along chemical gradients.

#### Cell mechanics and motion

We model cell mechanics and motion as in our prior work [14]: we update each cell’s position x_*i*_ by calculating its current velocity v_*i*_ based upon the balance of forces acting upon it. The main forces include cell motility, drag-like forces, and cell-cell and cell-matrix interaction forces: adhesion and “repulsion” (resistance to deformation and/or volume exclusion [26]). As in prior cell-centered models [14, 27,28], we apply an inertialess assumption 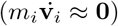 to explicitly solve for each cell’s velocity. As before [14], we model adhesion and repulsion with interaction potentials that depend upon each cell’s size, maximum adhesion distance, adhesion and repulsion parameters, and distance to other cells. The cell’s velocity v_*i*_ is given by

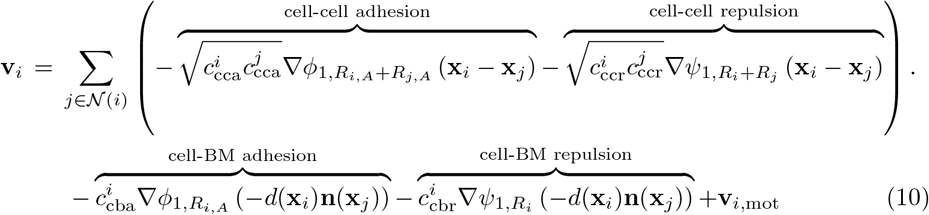

where as in [14] *φ_nR_* (x) is an adhesion interaction potential function which is zero for ||x|| > *R*, and approaches zero with smoothness given by *n*. Similarly, *ψ_n,R_*(x) is a repulsion interaction potential function that is zero for ||x|| > *R*. Thus, cell-cell mechanical interactions occur over finite distances. Here, 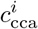 and 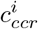 are the *i*^th^ cell’s cell-cell adhesion and repulsion parameters, *R_i_* is its radius, and *R_i,A_* is its maximum adhesion distance (typically a fixed multiple of *R_i_*). Note that if cell *i* and *j* have identical cell-cell adhesion and repulsion parameters *c*_cca_ and *c*_ccr_, then the cell-cell interaction coefficients simplify to the form we published in [14]; otherwise, this is a phenomenologic form that allows either cell’s coefficient to affect the cell-cell interaction strength, with equal and opposite effect on both cells. We plan future improvements to move beyond the current phenomenologic forms to more mechanistic models.

Also *d*(x) is the distance to the nearest basement membrane (if any), n(x) is a unit vector normal to the basement membrane, and so –*d*(x_*i*_)n (x_*i*_) points from the cell’s position x_*i*_ to the nearest point on the basement membrane, located at x_*i*_ – *d* (x_*i*_) n (x_*i*_). (See [29–31] for more information on level set (distance function) representations of surfaces, there applied to tumor growth models.) 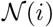 is the (finite) list of cells that could potentially interact with cell *i*. (See Key code optimizations.) Further references and the full forms of the potential functions are given in S1 Text.

The cell’s position is updated using the second-order Adam’s Bashforth discretization:

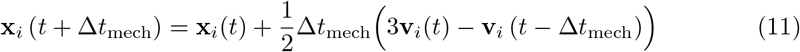

#### Cell cycling

PhysiCell includes a cell cycle modeling framework, where each cell cycle model is a collection of phases {*X_i_*}, transition rates {*r_ij_*} between the phases, and a cell division phase transition. As of Version 1.2.0, users can also set phase entry and exit functions (associated with the phases *X_i_*) that are executed at entry into or exit from the phase; these can be used to model processes such as mutation of cell parameters. The framework also allows users to set arrest functions (associated with the transition rates *r_ij_*) that block the transition. This is useful for modeling effects like volume restrictions. See the User Guide (S2 Text) for full details. As in [14], we use the phase transition rates to calculate the phase change probabilities in any time interval [*t,t* + Δ*t*]: the probability of transitioning from phase *X_i_* to phase *X_j_* in this time interval is given by

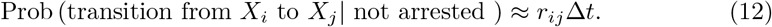

Users can set individual transitions *r_ij_* to have deterministic duration, with duration 1/*r_ij_*. See the User Guide (S2 Text) for full details.

Each cell agent tracks its current cell cycle phase 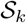 and its total time spent in that phase (*t_k_*). Users can change the transition rates at any time, in part based upon microenvironmental conditions (e.g., based upon oxygenation or cell contact).

As a concrete example, consider the “Ki67 Advanced” model from our prior work calibrating oxygen-dependent growth to Ki67 data in ductal carcinoma in situ (DCIS) [14, 15,32]. The phases are *K*_1_ (Ki67+ cycling cells, prior to cell division), *K*_2_ (Ki67+ cycling cells, after cell division), and *Q* (Ki67-quiescent cells). *K*_1_ and *K*_2_ have stochastic durations (with means *T*_1_ and *T*_2_). We model the transition rate from *Q* to *K*_1_ as

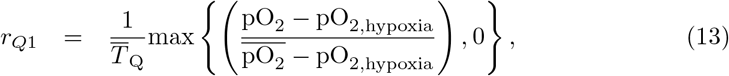

where cells spend a mean time of 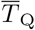 in the *Q* phase when 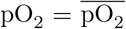. Cells double 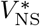 when transitioning from *Q* to *K*_1_ (to double their nuclear content), and they halve 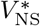 (and all the sub-volumes) when dividing into two daughter cells at the *K*_1_ → *K*_2_ transition. The full set of supported cell cycle models—along with reference parameter values—is given in S1 Text.

#### Cell death

PhysiCell currently includes models for two types of cell death: apoptosis (programmed cell death) and necrosis (unprogrammed cell death) [33]. At any time, each agent (with index *i*) has two death rates (*r_A,i_* for apoptosis, and *r_N,i_* for necrosis), which can be continually updated. For any death rate *r_i_* and any time interval [*t,t* + Δ*t*], the cell has a probability of entering the corresponding death state *D_i_*:

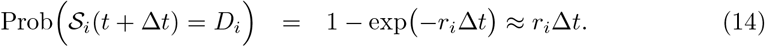

##### Apoptosis

Upon entering the apoptotic state, we set *f*_CN_ = 0 (to simulate shrinking and blebbing of the cytoplasm), 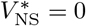 (to simulate degradation of the nucleus), and *f*_F_ = 0 (to simulate the active elimination of water from the cell). The rates *r*_N_, *r*_F_, and *r*_C_ are set to match time scales of cell volume loss in apoptotic cells. The cell is removed once its volume drops below a user-set threshold, or after mean duration of *T*_A_.

##### Necrosis

When a cell becomes necrotic, we set 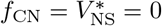 to model cytoplasmic and nuclear degradation. Early necrotic cells undergo oncosis (cell death-related swelling); we model this by setting *f*_F_ = 1. (Note that some regard oncosis as the actual death process, and necrosis as post-mortem cell degradation [34,35].) Once the cell volume passes a critical threshold, it lyses, and we set *f*_F_ = 0. The rate parameters *r*_F_, *r*_N_, and *r*_C_ are set to match expected time scales throughout necrosis [33]. PhysiCell includes codes to trigger necrosis deterministically or stochastically:

##### Deterministic Necrosis

This implements a common model of necrosis (see the review [2]), where cells instantly become necrotic whenever oxygenation pO_2_ drops below a threshold value pO_2_ threshold, as in our earlier work [14]. This is equivalent to the letting *r*_N_ → ∈.

##### Stochastic Necrosis

This model updates our prior work [14], based upon in *vitro* observations that cells can survive low oxygen conditions for hours or days. Here,

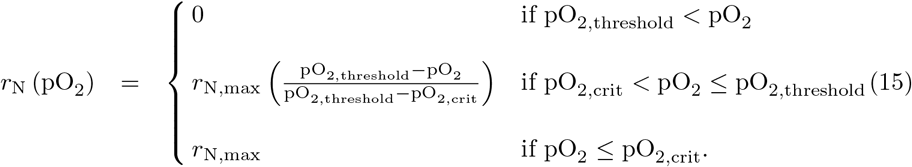

That is, necrotic death begins when pO_2_ < pO_2_ threshold, and the death rate ramps linearly until saturating at a maximum rate *r*_N,max_ for pO_2_ < pO_2_ crit. Equivalently, cells survive on average 1/*r*_N,max_ time in very low oxygen conditions [14].

### Numerical implementation

#### Time steps

PhysiCell has three time steps to model (fast) diffusive biotransport processes (Δ*t*_diff_; default 0.01 min), cell movement (Δ*t*_mech_; default 0.1 min), and (relatively slow) cell processes (Δ*t*_cells_; default 6 min). We use these time steps to set how frequently biotransport processes, cell movement processes, and cell phenotype processes are updated. See Fig. 1.

**Figure 1.**
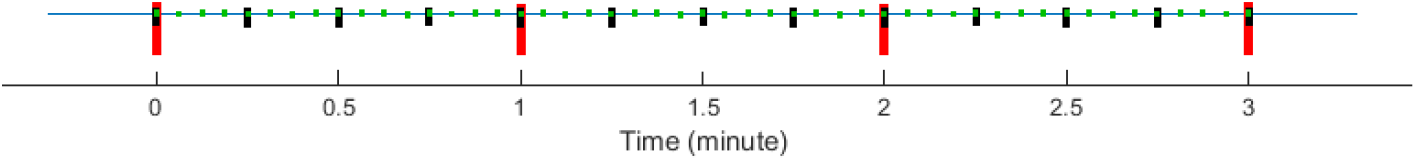
PhysiCell and multiple time scales: PhysiCell uses BioFVM to update the microenvironment at the short green tick marks, corresponding to Δ*t*_diff_. It updates cell mechanics (including cell position) less frequently at the medium black tick marks (Δ*t*_mech_), and it runs the cell volume and cycle/death models least frequently at the long red tick marks (Δ*t*_cen_). Note that the time steps shown are for illustrative purpose; the default step sizes are given in Time steps.

The default Δ*t*_diff_ was chosen for diffusion, decay, and uptake/secretion parameter values typical for the cancer and tissue engineering problems that drove PhysiCell’s development. In prior testing, relative errors did not exceed 5% for this value [6]. In mechanical relaxation tests for overlapping cells and compressed tumor spheroids, we found that Δ*t*_mech_ = 0.1 min gave solutions that converged at first-to-second order accuracy, had relative errors 5% or less, and avoided spurious oscillations and other artifacts for cell velocities under ~ 1 *μ*m/min (typical for cancer biology problems); see S1 Text. The cell cycle, death, and volume change models were numerically stable and first-order accurate with relative errors of 5% or less for Δtcells = 6 min. See S1 Text. Users should reduce Δ*t*_cells_ for problems with faster phenotypic processes. Users anticipating faster cell movement (e.g., motile bacteria) should reduce Δ*t*_mech_. We recommend setting 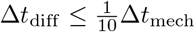.

Mathematically, this time scale separation allows us to hold cell positions fixed (quasi-static) when updating the PDE solutions, and then hold the chemical fields fixed when updating cell positions and phenotypes. We have used similar techniques in nonlinear continuum models of tumor growth (slow time scale) in heterogeneous biochemical microenvironments (fast time scale) [29–31].

#### Estimated computational cost scaling

We now assess the computational effort needed for each iteration in the main program loop. (See Overall program flow.) Step 1 (save simulation data), Step 3 (run cell processes), and Step 4b (update positions) clearly entail a constant amount of work for each cell. Thus, summing these steps over all cells *n*(*t*) requires 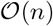 work. By prior analysis, BioFVM (Step 2) also scales linearly in *n*(*t*) [6].

Step 4a (update velocities) is the most computationally expensive step. In straightforward implementations, each cell tests for mechanical interaction with *n* – 1 other cells, giving an 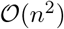 total computational cost at each time step. However, the interaction data structure (see Key code optimizations) restricts interaction testing to a smaller set 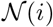.

In Expanded computational cost estimates, we show that each 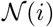 has at most *N*_max_ cells. Thus, Step 4b has a fixed maximum cost for each cell, and the cost of the loop scales linearly in *n*.

#### Key code optimizations

To prevent computational costs from scaling quadratically in the number of cells, we designed a cell-cell interaction data structure (IDS) that efficiently estimates a set 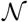 of possible neighbor cells for each cell agent. See S1 Text for further detail.

PhysiCell uses OpenMP to parallelize most loops over the list of cells. This includes sampling the microenvironment, updating cell phenotype parameters, advancing the cell cycle or death model, advancing the volume model, running any custom function, and calculating the cell velocity. We do not parallelize loops that change the IDS: cell division, cell removal, and updating the cell position.

As discussed above, we defined three separate computational step sizes (Δ*t*_diff_ < Δ*t*_mech_ < Δ*t*_cells_) to take advantage of the multiple time scales of the multicellular system. As indicated in the overall program flow above, we update each process according to its own time step, rather than at each simulation step. Fig. 1 further illustrates how the multiple times steps reduce the computational cost. See S1 Text for further detail and the default step sizes for cancer biology and tissue engineering.

#### Expanded computational cost estimates

Most of the simulation steps have computational cost that scales linearly in the number of cells. (See Estimated computational cost scaling.) The step that requires additional analysis (and relies upon PhysiCell’s IDS) is the step where cell-cell mechanical interactions are used to set the cell velocities. Bounding this computational costs requires that we find a fixed upper bound on the number of cell-cell interactions, so that the computational cost is 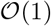 for single cells, and 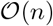 for all the cells.

We estimate an upper bound on the of cells in any voxel *B_i_* by

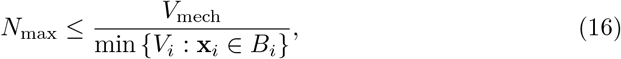

where *V*_mech_ is the fixed volume of the voxels in the interaction testing data structure. For cycling cells with “mature” volume *V*, we have 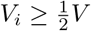. By default, dead cells are removed when 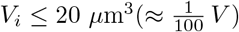. Since a typical 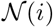 is constructed from up to 27 such voxels, we have

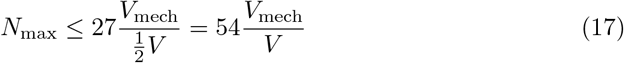

for simulations dominated by live cells, and

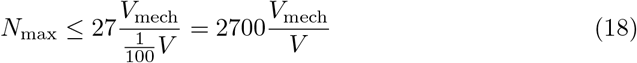

for simulations dominated by dead cells. Thus, the computational cost for a single cell’s mechanical interactions is bounded by a fixed constant, and the total cost over all cell-cell mechanical interactions scales linearly in *n*. The slope of the cost-versus-n curve may be shallower for early, non-necrotic simulations, and it can be up to a factor of 100 steeper for necrosis-dominated simulations. In some cases, simulations may temporarily show a nonlinear relationship with n when transitioning from non-necrotic to necrotic.

### Convergence and validation testing

We performed convergence testing on all the major components of PhysiCell. BioFVM was previously tested as first-order accurate in Δ*t*, second-order accurate in Δ*x*, and sufficiently accurate at Δ*x* = 20 *μ*m and Δ*t*_diff_ = 0.01 to 0.05 min for tumor growth problems [6]. We performed two tests for cell-cell mechanics and motion: First, we placed two cells in partial overlap, simulated their relaxation to equilibrium, and measured the cell spacing at several times. Second, we created a compressed cluster of 50,000 cells, simulated its mechanical relaxation to equilibrium, and measured its diameter at several times. Both tests converged to first-order accuracy in Δ*t* at all measured times, showing that PhysiCell converged in both short-time mechanical dynamics and in long-time behavior. Δ*t*_mech_ ~ 0.1 min gives sufficient accuracy for typical cancer problems.

We simulated the volume model for a single proliferating, apoptotic, and necrotic cell, and measured the sub-volumes at multiple times. It converged with first-order accuracy in Δt at all tested times, and Δ*t*_cell_ = 6 min gave sufficient accuracy. We tested the stochastic transition codes by simulating the Ki67-advanced cell cycle model and the apoptosis death model (with stochastic duration), and measuring the sub-population counts and population fractions over time for several values of Δ*t*_cell_. For each Δ*t*, we performed 100 simulations and compared the mean solution behavior against known coarse-grained ODE model behavior. Δ*t*_cell_ = 6 min and 60 min both gave an excellent match between the PhysiCell behavior and theory for all the compared curves. See S1 Text for full testing results.

### Performance testing (summary)

By our testing, recent quad-core desktop workstations (with hyperthreading, for 8 total execution threads) can simulate 10-30 days in systems of up to 10^5^ to 10^6^ cells in 3 days or less (wall time). Single HPC compute nodes (typically two 6-8 core processors, with hyperthreading and 24-32 execution threads) can simulate larger systems up to ~2 million cells in about 2 days. Future releases of PhysiCell will address current performance bottlenecks; see Availability and Future Directions. The Results will give a demonstration of 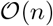 computational cost scaling.

## Results

We demonstrated PhysiCell’s potential to simulate large multicellular systems—and its ability to test the emergent tissue-scale effects of cell-scale hypotheses—on several examples arising from cancer biology and synthetic multicellular systems bioengineering. For the first two examples, we compared the impact of the deterministic and stochastic necrosis models. (See Cell death above.) We used the Ki67-advanced cell cycle model with deterministic *K*_1_, *K*_2_, and A phase durations for the first two examples. (See Cell cycling.) We used a simpler “live cells” cycle model [22] in the remaining examples, where live cells proliferate with a variable birthrate, apoptose, or necrose. We provide detailed parameter values in S1 Text for the HDS and DCIS examples, and the full source code and postprocessing routines for both examples in every PhysiCell download. Full source code and simple build instructions are provided for the remaining examples. Reference simulation outputs for the first two examples are available at http://PhysiCell.MathCancer.org.

### Test platforms

The Hanging drop tumor spheroids and Ductal carcinoma in situ (DCIS) examples were tested on (1) a desktop workstation (quad-core Intel i7-4790, 3.60 GHz, 8 execution threads, 16 GB memory) with mingw-w64 (`g++` ver. 4.9.1) on 64-bit Windows 7, and (2) a single HPC compute node (dual 6-core Intel Xeon X5690, 3.47 GHz, 24 execution threads, 48 GB memory) with `g++` (ver. 4.8.4) on Ubuntu 14.04. The tests were performed using PhysiCell 1.0.0, although release 1.2.0 has updated the tests for compatibility. The CPU architecture was newer on the desktop (2014 Haswell) than on the HPC node (2011 Westmere). The newer “Biorobots”, Anti-cancer biorobots, Cancer heterogeneity and immune response, and Adding an immune response examples were tested on a quad-core Intel i7-4770K, 4.06 GHz, 8 execution threads, 32 GB memory, using PhysiCell Version 1.2.1 with g++ 7.1.0 on 64-bit Windows 10 (via MinGW-w64).

### Hanging drop tumor spheroids

Hanging drop spheroids (HDS)—a 3-D cell culture model where a small cluster or aggregate of tumor cells is suspended in a drop of growth medium by surface tension—are increasingly used to approximate 3-D *in vivo* growth conditions [36]. Unlike traditional 2-D monolayer experiments, HDSs allow scientists to investigate the impact of substrate gradients on tumor growth, particularly oxygen gradients. Their relatively simple geometry makes them ideal for testing computational models.

We simulated HDS growth by placing an initial cluster of ~ 2300 cells in an 8 mm^3^ fluid domain, with Dirichlet conditions pO_2_ = 38 mmHg (5% oxygen: physioxic conditions [37]) on the computational boundary. The simulation results are shown in Fig. 2 for deterministic necrosis (left column) and stochastic necrosis (right column), at 4, 8, and 16 days. In Fig. 3, we show the tumor diameter (left panel) and number of agents (right panel) versus time. Both simulations reached ~ 10^6^ cells by 18 days. See the simulation videos S1 Video and S2 Video.

**Figure 2.**
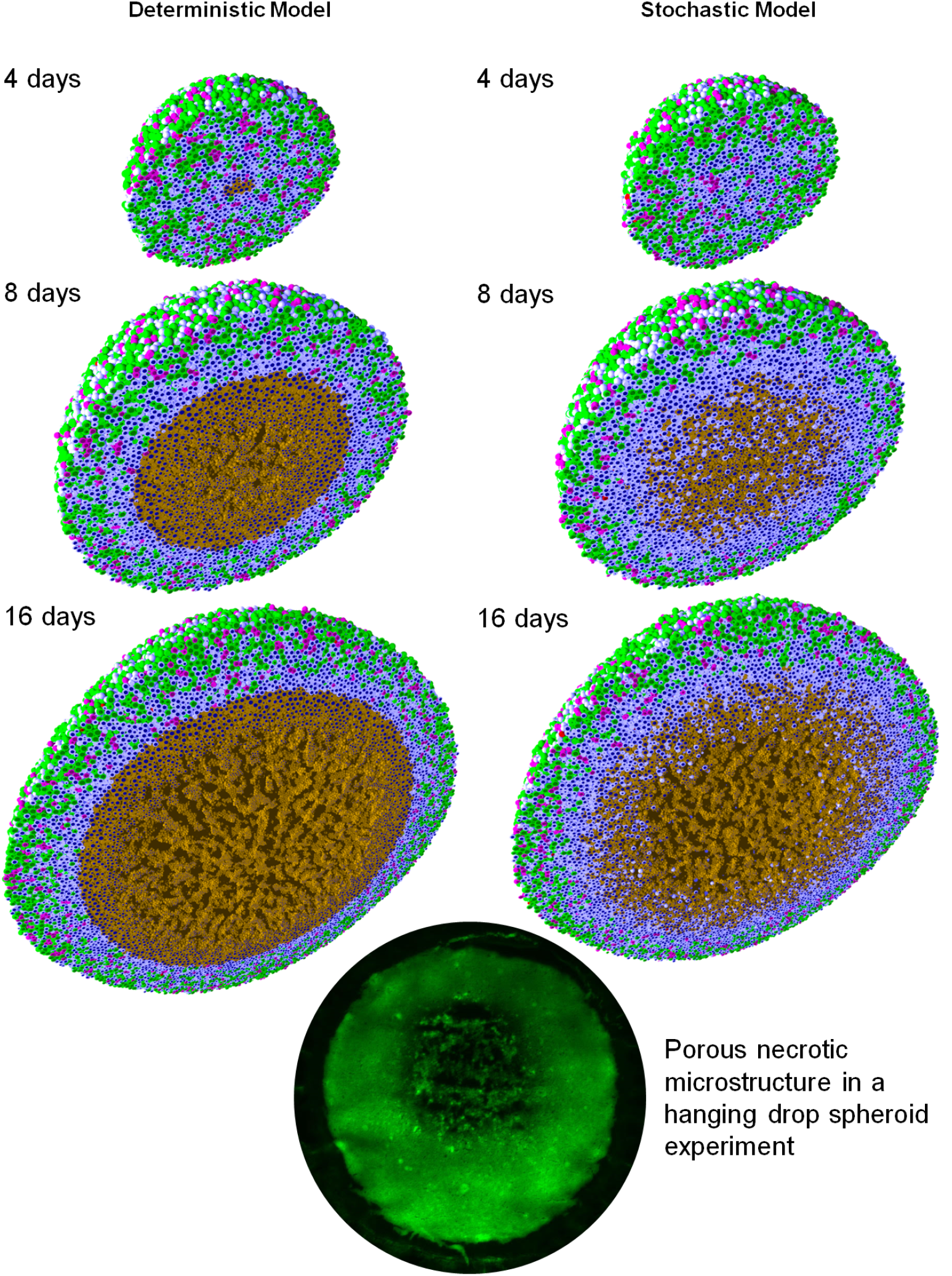
**Hanging drop spheroid (HDS) simulations** with deterministic necrosis (left) and stochastic necrosis (right), plotted at 4, 8, and 16 days. Videos are available at S1 Video and S2 Video. ***Legend***: Ki67+ cells are green before mitosis (*K*_1_) and magenta afterwards (*K*_2_). Pale blue cells are Ki67-(Q), dead cells are red (apoptotic) and brown (necrotic), and nuclei are dark blue. **Bottom**: Hanging drop spheroid experiment (HCC827 non-small cell lung carcinoma) showing a similar necrotic core microstructure. PhysiCell is the first simulation to predict this structure arising from cell-scale mechanical interactions. Image courtesy Mumenthaler lab, Lawrence J. Ellison Center for Transformative Medicine, University of Southern California.

**Figure 3.**
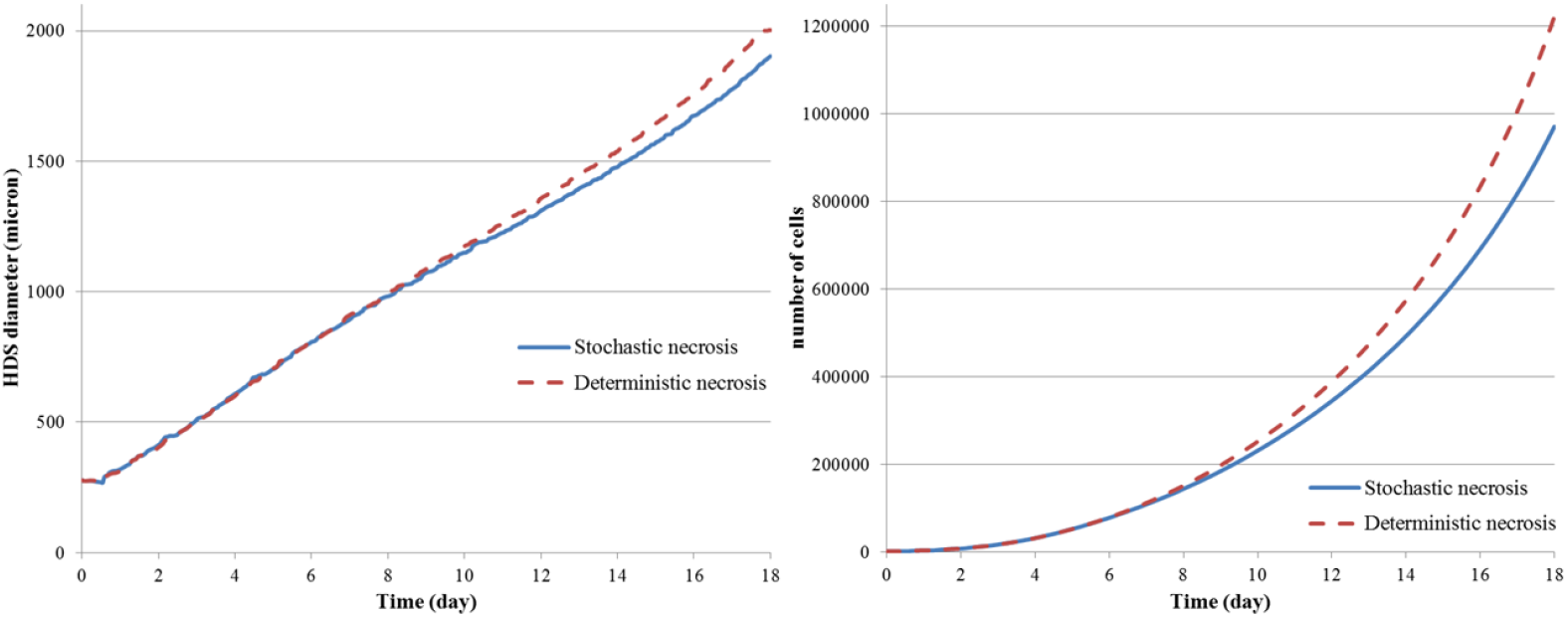
HDS growth: *Left:* The deterministic and stochastic necrosis models both give approximately linear growth (left), but the HDS with deterministic necrosis model grows faster (~ 5% difference in diameter at day 18). ***Right***: The HDS with stochastic necrosis has fewer cells than the deterministic model (~ 26% difference in cell count at day 18), due to its delay in necrosis. The difference in cell count is larger than the difference in tumor diameter because most of the difference lies in the number of necrotic cells, and necrotic cells are smaller than viable cells.

#### Deterministic versus stochastic necrosis

Both models yielded similar dynamics. Hypoxic gradients emerged quickly, limiting (pO_2_-dependent) cell division to the outermost portions of the tumors. This, in turn, lead the tumor diameters to grow linearly (at similar rates); see Fig. 3. This matches our theoretical expectations for a spheroid of radius *R*(*t*) whose growth is restricted to an outer layer of fixed thickness *T*:

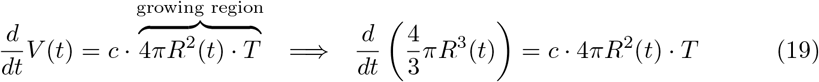

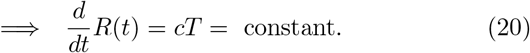

In both models, the innermost portion of the necrotic core developed a network of fluid voids or cracks. This phenomenon emerges from competing biophysical effects of the multicellular system and its cell-scale mechanical details: necrotic cells lose volume, even as they continue to adhere, leading to the formation of cracks. To our knowledge, this is the only model that has predicted this necrotic tumor microarchitecture, which would be very difficult to simulate by continuum methods except with very high-resolution meshes comparable to the ~ 1 to 10 *μ*m feature size. These cracked necrotic core structures have been observed with *in vitro* hanging drop spheroids (e.g., [5, 38,39]) See Fig. 2.

There were notable differences between the models. The deterministic model had a sharp perinecrotic boundary between the viable and necrotic tissues, whereas the stochastic model demonstrated a perinecrotic transition zone with substantial mixing of viable and necrotic cells. Because cells do not immediately necrose in the stochastic model, it retained a center of quiescent viable cells longer than the deterministic model. The growth curves for the deterministic and stochastic models appear to diverge after approximately 8 days, when the deterministic necrotic core is better defined with more cracks than the stochastic core. This may be due to differences in hypoxic gradients (the tumor with more void spaces will have shallower oxygen gradients, and hence more cell cycle entry), but further simulations would be required to rule out stochastic effects. Interestingly, the stochastic model’s growth curve appears to run parallel to the deterministic curve for later times, once its necrotic core becomes better defined.

#### Performance scaling

Throughout the simulations, the computational cost (the wall time required to simulate one hour) scaled approximately linearly with the number of agents present in the simulation, on both the desktop workstation and the HPC node; see Fig. 4. (See also Estimated computational cost scaling.) Increasing the number of execution threads improved performance, even when running on slower processor cores. See the right panel in Fig. 4, where moving from the newer 8-threaded machine to the older 24-threaded machine improved performance by a factor of 2 to 2.5.

**Figure 4.**
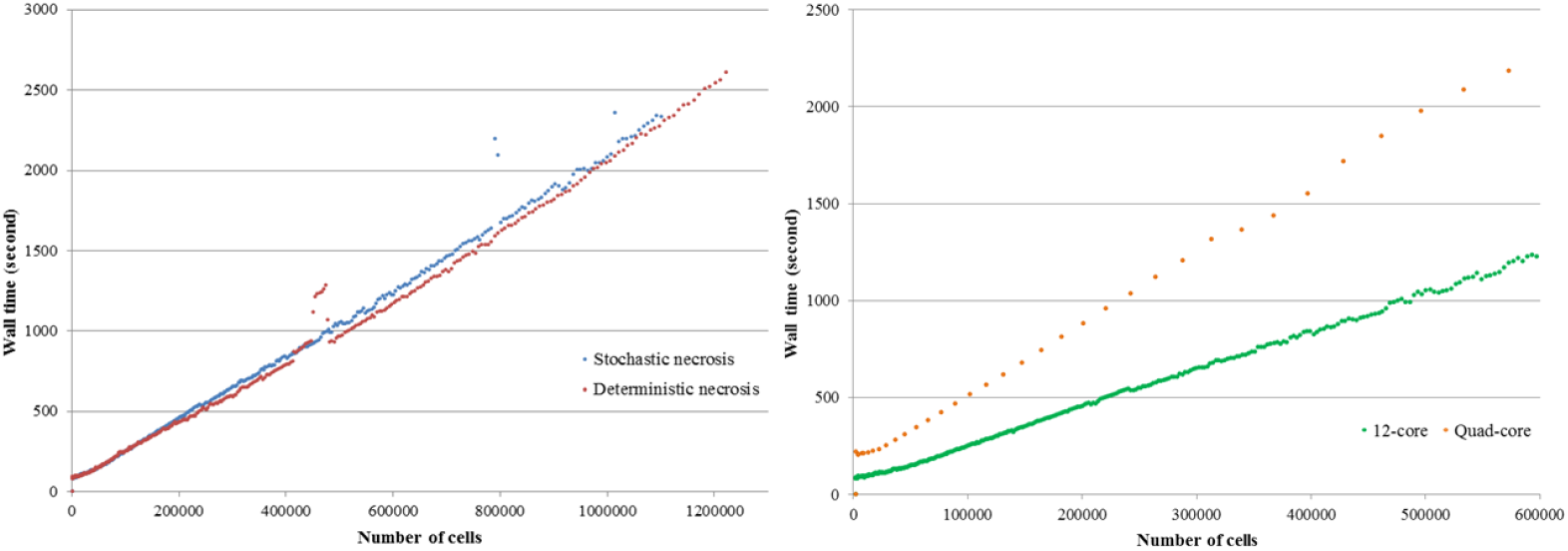
HDS computational cost scaling: ***Left***: Wall-time vs. cell count for the stochastic (red) and deterministic (blue) necrosis necrosis models on a single HPC compute node. Both models show approximately linear cost scaling with the number of cell agents. ***right***: Wall time vs. cell count for stochastic necrosis model on the desktop workstation (orange) and the single HPC node (green).

The simulations reached ~ 10^6^ cells on our HPC tests after 67 hours (deterministic, 17 simulated days) to 76 hours (stochastic, 18.2 simulated days) of wall time, including saving full simulation output data once per simulated hour. See Fig. 3. The desktop workstation simulated past 573,000 cells (about 14.6 days of simulated time) in approximately 80 hours of wall time. The desktop tests did not run out of memory, and the simulations can be completed to the full 18 days and 10^6^ cells if needed.

### Ductal carcinoma in situ (DCIS)

DCIS is a pre-malignant breast condition where epithelial cells (“tumor cells”) divide abnormally to fill the breast duct lumen. Oxygen can only reach the tumor cells by diffusion from outside the duct, leading to the emergence of hypoxia and an inner necrotic core. See [14, 15, 32] for further biological and clinical discussion. As in [14], we approximate a partly-filled breast duct as a 3-D “test tube” with a level set function representation. Cells adhere to cells and the duct wall; cells and the duct wall push against cells to resist deformation. Oxygen diffuses from the duct wall and is consumed by tumor cells. The rate of cycle entry increases linearly with pO_2_ (see Cell cycling).

In Fig. 5, we show DCIS simulations in a 1 mm segment of breast duct (317.5 *μ*m diameter), using deterministic necrosis (left side) and stochastic necrosis (right side), plotted at 10 and 30 days. See also S3 Video and S4 Video. As in prior work [14], the simulations predict cell-scale details observed in DCIS pathology, such as the appearance of pairs of Ki67+ daughter cells, the spatially isolated apoptotic cells (which arises from the model assumption that apoptosis is a stochastic, low-frequency event that is independent of oxygenation), and the higher occurrence of Ki67+ cells near the duct wall (where we modeled the probability of cell cycle entry as proportional to oxygenation).

**Figure 5.**
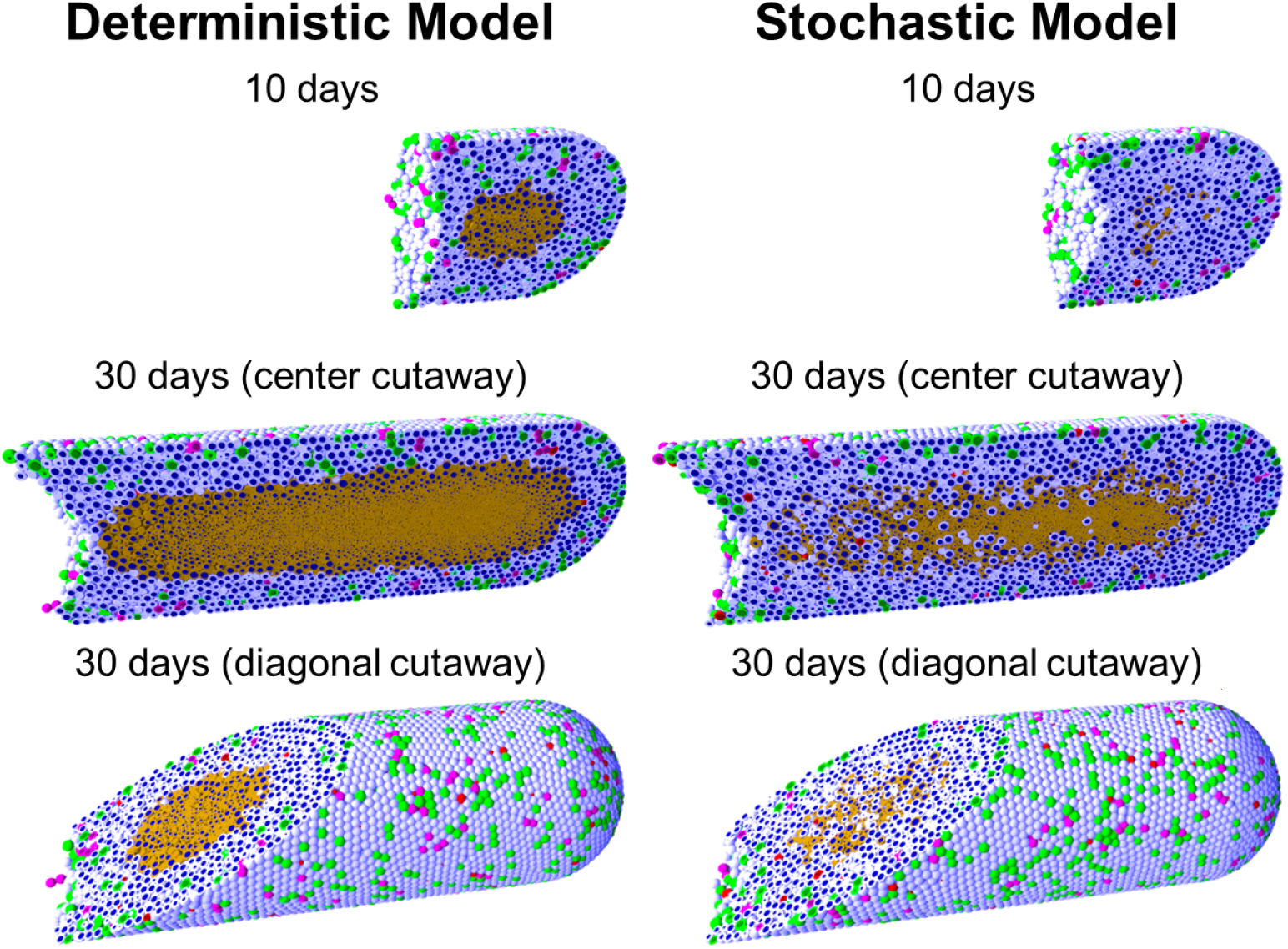
**Ductal carcinoma in situ (DCIS) simulations** with deterministic necrosis (left) and stochastic necrosis (right), plotted at 10 and 30 days (multiple views). Videos are available at S3 Video and S4 Video. The figure legend is the same as Fig. 2.

#### Comparison of necrosis models; comparison with the spheroid example

As in the HDS example, the deterministic model had a sharp, smooth perinecrotic boundary, whereas the stochastic model demonstrated a perinecrotic boundary region with mixed viable and necrotic cells. In the stochastic model, proliferation halted in the duct center, but necrosis appeared later. The perinecrotic mixing effect was most pronounced at the leading edge of the tumor, where tissue was transitioning from non-hypoxic/non-necrotic to necrotic. Areas with longer-term hypoxia had smoother necrotic boundaries. This effect did not emerge in the HDS example due to its symmetry.

Interestingly, the mechanical “cracks” seen in the tumor spheroids do not appear here, because the breast duct compresses the necrotic core to collapse any fluid-filled voids. This shows the importance of the 3-D geometry and the biophysical impact of the basement membrane, as well as the need to account for such effects when approximating *in vivo* conditions with bioengineered model systems.

Both models gave approximately the same growth rate of ~ 1 cm/year (Fig. 6, left). We cannot select one model over the other based solely upon continuum-scale, coarsegrained outputs. However, we could further assess the models by comparing their distinct differences in multicellular-scale patterning to DCIS pathology. This further highlights the need and potential for multicellular modeling in evaluating cell-scale hypotheses.

**Figure 6.**
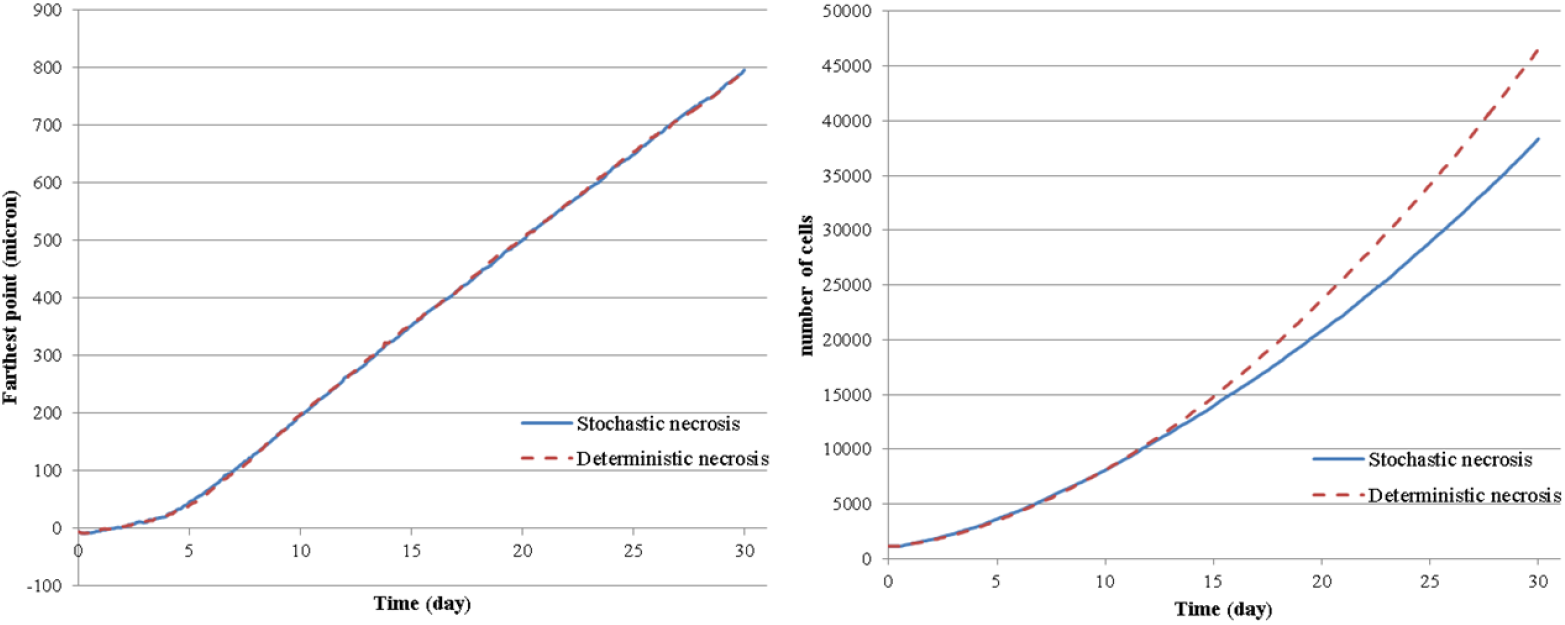
DCIS growth: The deterministic and stochastic necrosis models both result in linear DCIS growth at approximately 1 cm/year (left), even while their cell counts differ by 21% by the end of the simulations (right).

#### Comparison with prior 2-D modeling results

In 3D, neither necrosis model reproduced the mechanical “tears” between the proliferative rim and the necrotic core predicted by earlier 2-D simulations [14]; this is because more viable tissue is fluxing into smaller necrotic areas in the 3-D geometry compared to the 2-D geometry.

### Additional PhysiCell examples

In the following examples, we demonstrate PhysiCell in applications involving interactions via contact and chemical factors between multiple cell types, in 2-D and 3-D simulations. Most of the examples include cell motility, “custom” phenotype rules, additional “custom” mechanics, and custom data.

Please visit http://www.mathcancer.org/blog/physicell-sample-projects/ for instructions on building, running, and visualizing the following sample projects. The sample projects are structured to mirror the intended design of user projects: a project-tailored `Makefile` and `main.cpp` in the main directory, and most custom code in the project-tailored functions in the `custom_modules` directory.

The instructions can also be found in the User Guide (S2 Text, found in the documentation folder of every PhysiCell download) and the Quickstart guide (S3 Text, found in the root directory of every PhysiCell download).

#### “Biorobots”

First, we tested PhysiCell for its potential in aiding in the design of synthetic multicellular systems. We investigated rulesets to create a cellular cargo delivery system, including the following main components:

1. “Director” cells secrete a diffusible chemoattractant *c*_Ţ_ and are otherwise static. They use the typical cell-cell repulsion mechanics.
2. “Cargo” cells have a surface receptor 0 ≤ *R* ≤ 1. When *R* =1, they secrete a diffusible chemoattractant c2. When R = 0, they stop secreting the factor. They use the typical cell-cell repulsion mechanics.
3. “Worker” cells can either be adhered or not adhered to cargo cells.

(a) When they are unadhered, they perform biased random migration towards gradients of ∇_c_2__. Whenever they touch a cell, they test for presence of receptor *R*. If *R* is expressed (i.e., the worker has found cargo), the worker forms an elastic adhesion with the cell and sets *R* = 0 on the cargo cell.
(b) When they are adhered, they perform biased random migration towards ∇_c_1__. When *c*_1_ exceeds a threshold, they break their adhesive bond.

As a simple model of this targeted cell adhesion, we used a custom_celljrule to implement:

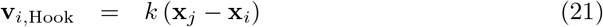

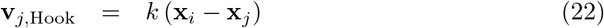

for adhered cells *j* and *i*. We then add these to the cell’s velocities. PhysiCell evaluates the custom rule when evaluating cell mechanics (velocity update).

Simulation outputs are shown in Fig. 7; see also S5 Video. As we can see, the cargo and worker cells successfully interact to modulate their behaviors. Notably, the worker cells are seen making multiple transits from the supply of cellular cargo to the directors, showing the robustness of both the rules and their implementation in PhysiCell.

**Figure 7.**
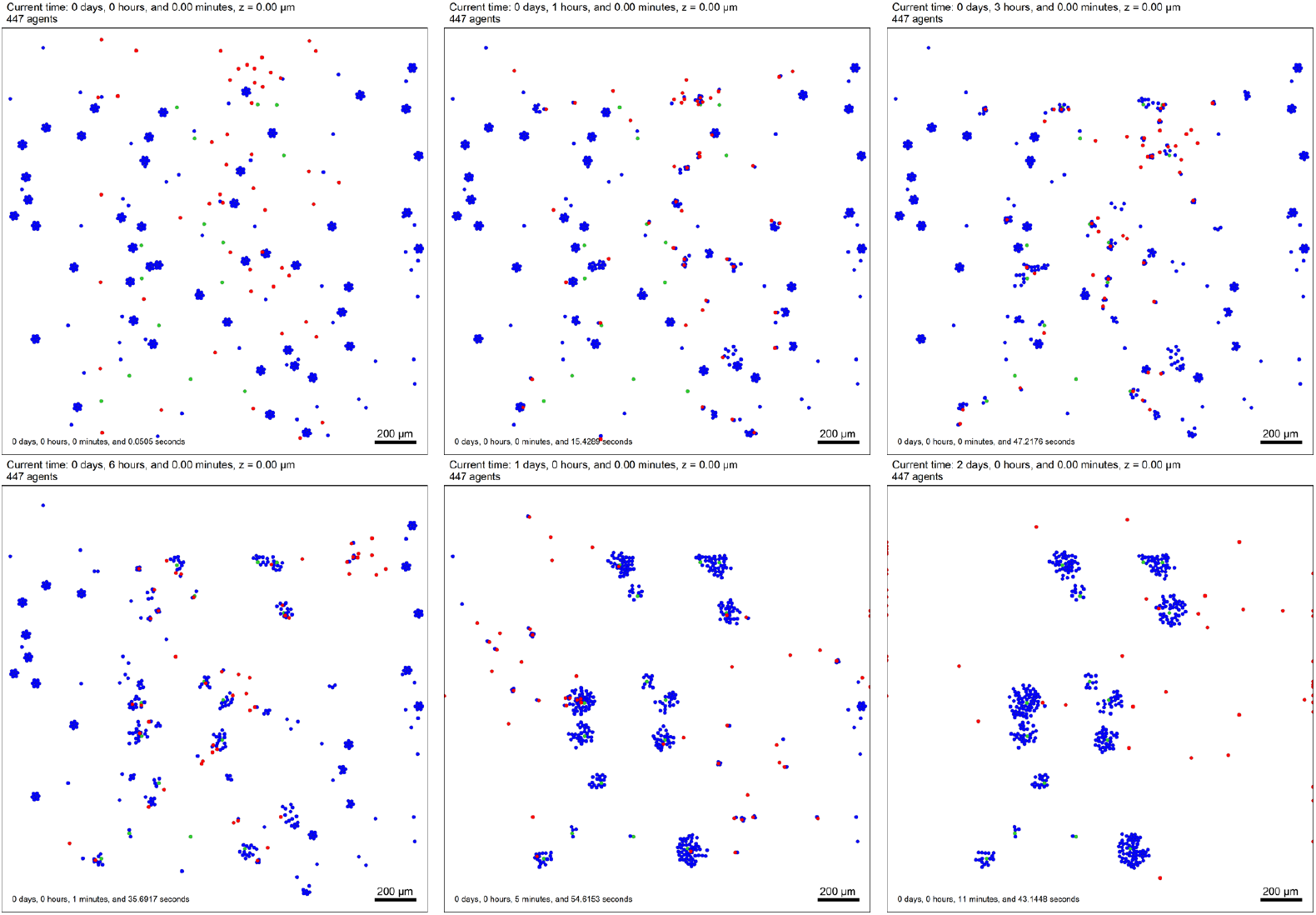
“Biorobots” example. Director cells (green) release a chemoattractant *c*_1_ to guide worker cells (red). Cargo cells (blue) release a separate chemoattractant *c*_2_. Unadhered worker cells chemotax towards ∇*c*_2_, test for contact with cargo cells, form adhesive bonds, and then pull them towards the directors by following ∇*c*_1_. If *c*_1_ exceeds a threshold, the worker cells release the cargo and return to seek more cargo cells, repeating the cycle. A video is available at S5 Video.

This and the following examples also demonstrate the intended arrangement of projects: users do not modify the contents of `./core/`, but instead place their codes in `./custom_modules/`, include these in a main project `main.cpp`, and modify the `Makefile` to compile and link the components. This design is intended to allow users to update the PhysiCell and BioFVM core components without overwriting their own customizations.

#### Anti-cancer biorobots

We adapted the “biorobots” to test their potential as an anti-cancer treatment. Many proposed cancer therapies attempt to target cancer cells by finding unique surface or other molecules to target, so that drugs can be conjugated to custom antibodies or encapsulated in custom nanoparticles. These still generally rely upon passive delivery of the therapeutics, even though cancers are often poorly perfused. See e.g. [40,41]. We tested the “biorobots” as a potential solution, with the following modifications:

1. There are no “director” cells. Instead, cancer cells consume oxygen (as in prior examples), which creates an oxygen gradient that can be leveraged for worker cell “homing.”
2. Adhered “cargo” cells detach themselves from “worker” cells when pO_2_ < pO_2_ drop and secrete a therapeutic compound [drug] that diffuses with the typical form given in Biochemical microenvironment.
3. Adhered worker cells perform biased random migration along −∇pO_2_.
4. In any time interval [*t,t* + Δ*t*_cells_], cancer cells accumulate drug-induced damage according to:

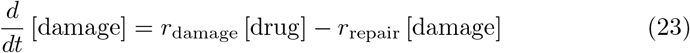

where *r*_damage_ and *r*_repair_ are the damage and repair rates. Each tumor cell agent tracks its own level of damage. In the same time interval the probability of a cell apoptosing due to the drug is

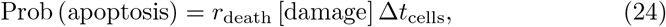

where *r*_death_ is the death rate when [damage] = 1.
5. Unadhered worker cells disabled motility if c_1_ falls below a threshold value.

We simulated 1 week of tumor growth, “injected” a mixture of 10% worker cells and 90% cargo cells near the tumor, and set the parameter pO_2_ drop to 10 mmHg. In the simulation (Fig. 8 and S6 Video), overall the cargo cells were delivered into the tumor (and they can be seen pushing tumor cells out of the way), tumor cells were indeed killed. However, once enough cancer cells were killed, hypoxia was reduced so that worker cells clustered near the oxygen minimum, but no longer released their “cargo” (because pO_2_ > pO_2_ drop throughout the domain). Setting pO_2 drop_ = 15 mmHg reduced but did not eliminate this behavior. See Fig. 8 and S6 Video. Thus, an “anti-cancer biorobot” system as explored here could potentially be beneficial (in particular, homing towards and penetrating tumors without need for cancer-specific targets), but the “cargo release” rules need to be carefully engineered. Such a system could potentially activate and deactivate to keep a tumor cell population in control, and to reduce hypoxia (which is known to drive cancer cell adaptation to more aggressive phenotypes [42,43]).

**Figure 8.**
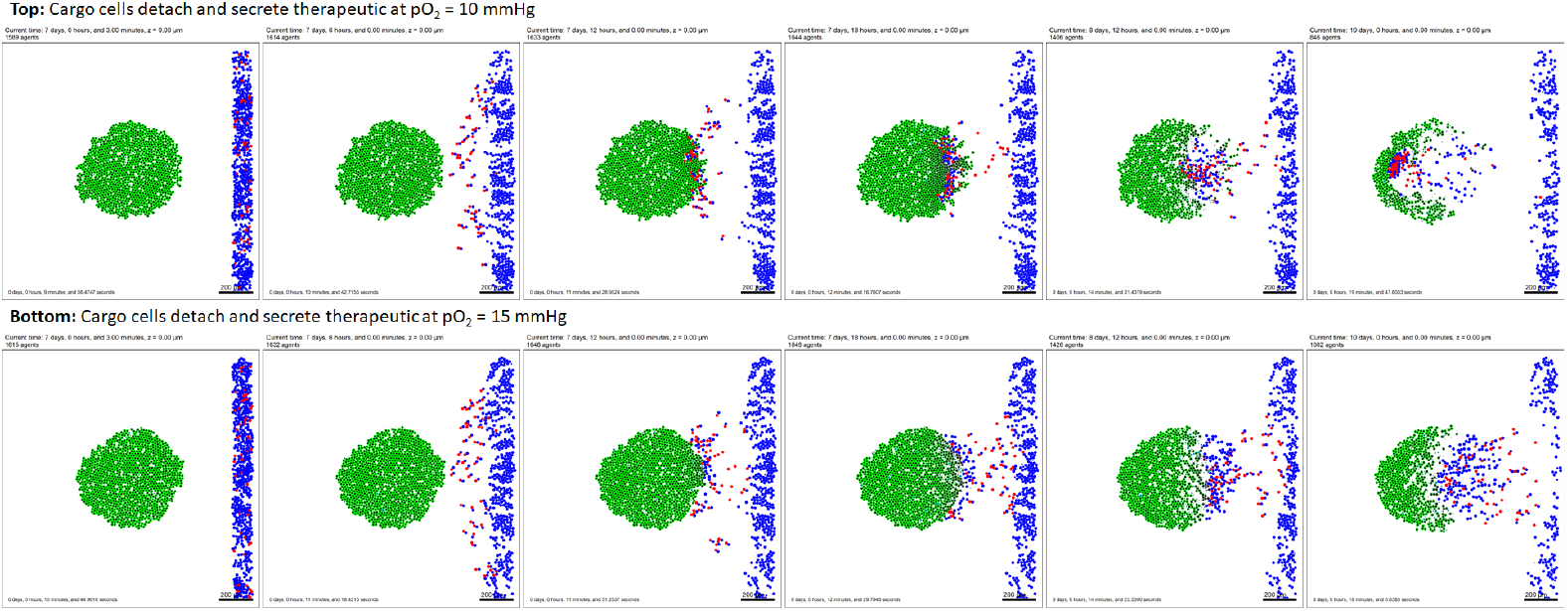
Anti-cancer “biorobots” example. By modifying the worker cells in the previous example (Fig. 7) to move up hypoxic gradients (along -∇pO_2_) and drop their cargo in hypoxic zones, we can deliver cargo to a growing tumor. In this example, the cargo cells secrete a therapeutic that induces apoptosis in nearby tumor cells, leading to partial tumor regression. A video is available at S6 Video.

#### Cancer heterogeneity and immune response

Next, we applied PhysiCell to another area of interest in the cancer community: heterogeneity. We seeded an initial tumor, and assigned each cell a random expression of a mutant “oncoprotein” 0 ≤ *p* ≤ 2. We started the simulation with a normal distribution of *p*, with mean 1 and standard deviation of approximately 0.3. We used the same oxygen-dependent stochastic cell cycle entry and necrosis as in prior examples. We modified the model to set the rate of cycle entry to scale proportionally to *p*, so that increased p increases the rate of cell division. The initial, intermediate, and final morphologies of the tumor are plotted in Fig. 9, along with the changing histograms of *p*. In the plot, cell color ranges from blue (*p* = 0) to yellow (*p* = 2). We can see clear selection for the yellower cells with greater expression of *p*. Moreover, see that while the initial tumor began with a uniform “salt and pepper” distribution of blue and yellow cells and a symmetric morphology, symmetry was broken by the end as regions with higher initial p and greater access to oxygen form dominant focal growths of “yellow” clones. This demonstrates further stochasticity in the simulation that would be difficult to predict with continuum approaches. See S7 Video to better examine these dynamics.

**Figure 9.**
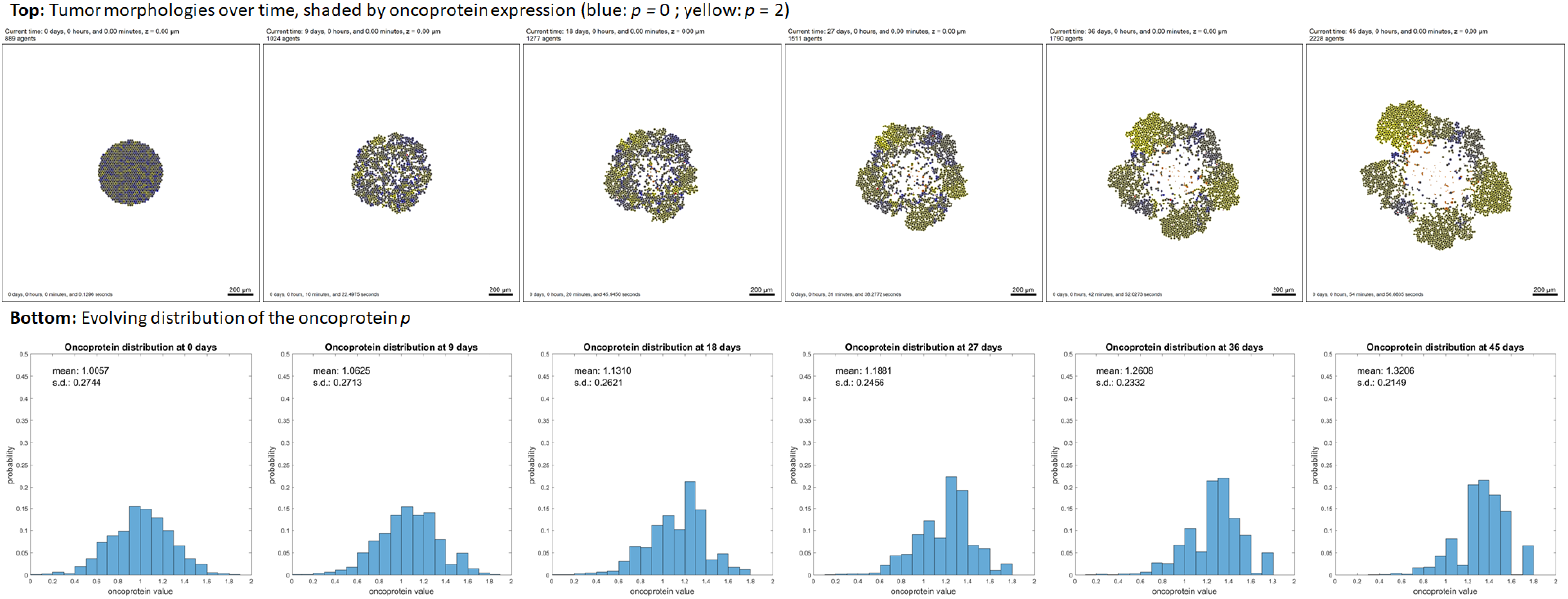
Cancer heterogeneity example. Each cell has an independent expression of a mutant “oncoprotein” *p* (dimensionless, bounded in [0,2]), which scales the oxygen-based rate of cell cycle entry. Blue cells have least *p*, and yellow cells have most. Initially, the population has normally distributed *p* with mean 1, standard deviation 0.3, and a “salt and pepper” mixed spatial arrangement. The proliferative advantage for cells with higher *p* leads to selection and enrichment of the most yellow cells. Stochastic effects lead to emergence of fast-growing foci and a loss of tumor symmetry. A video is available at S7 Video.

#### Adding an immune response

We extended the heterogeneity example to 3D, and integrated a simple model of an immune attack on the heterogeneous tumor:

1. All cancer cells consume oxygen as before. They also secrete an immunostimulatory factor *c* (e.g., a chemokine such as basic fibroblast growth factor (bFGF) [44]), which diffuses according to the standard PDE we introduced earlier.
2. As a simple model of immunogenicity, the mutant oncoprotein is assumed to increase immunogenicity proportionally to *p*, similarly to mutant tumor-associated epitopes being presented on MHCs (major histocompatibility complexes) [45,46].
3. Unadhered immune cells perform biased random migration (in our simulations *b* = 0.5) towards ∇_*c*_ and test for contact with cells. If they detect contact, they form an adhesion using the same model as in the “Biorobots” example and switch off motility. In any time interval [*t, t* + Δ*t*], we give the cell a probability of forming an adhesion regulated by

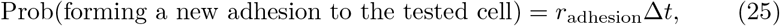

where *r*_adhesion_ is the immune cell’s rate of forming new cell adhesions.
4. While adhered, immune cells attempt to induce apoptosis in the adhered cell with probability

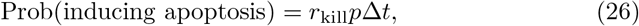

where *p* is used as our surrogate marker for immunogenicity (see above), and r_kill_ is the rate at which adhered immune cells kill tumor cells with *p* = 1. If they induce apoptosis, they detach and resume their search for new cells.
5. Adhered immune cells have a probability of detachment given by

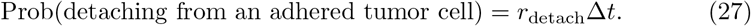

In Fig. 10 and S8 Video, we first simulate two weeks of tumor growth without an immune reaction. As in the prior example, we see the initially symmetric tumor develop asymmetric focal growths of “yellow” cells, again showing the selection for cells with the most oncoprotein. At two weeks, we introduce 7500 immune cells (red) that invade the tumor. Apoptotic tumor cells are labeled in cyan. The immune cells continue to migrate towards the center of the tumor up gradients of *c*, and within a few days, the tumor cell population is drastically reduced, with more “blue” cells (less immunogenic) remaining than “yellow cells” (most immunogenic). However, because the immune cell migration was strongly biased along ∇*c*, they pass by some tumor cells at the outer periphery. These surviving cells repopulate the tumor. This highlights the importance of stochasticity in immune cell migration; if homing is too strong, immune cells cannot mix with tumor cells, leading them to cluster in dense regions which can only interact with tumor cells on their edges. Less biased migration would increase mixing of cancer and immune cells and increase the efficacy of the immune attack.

**Figure 10.**
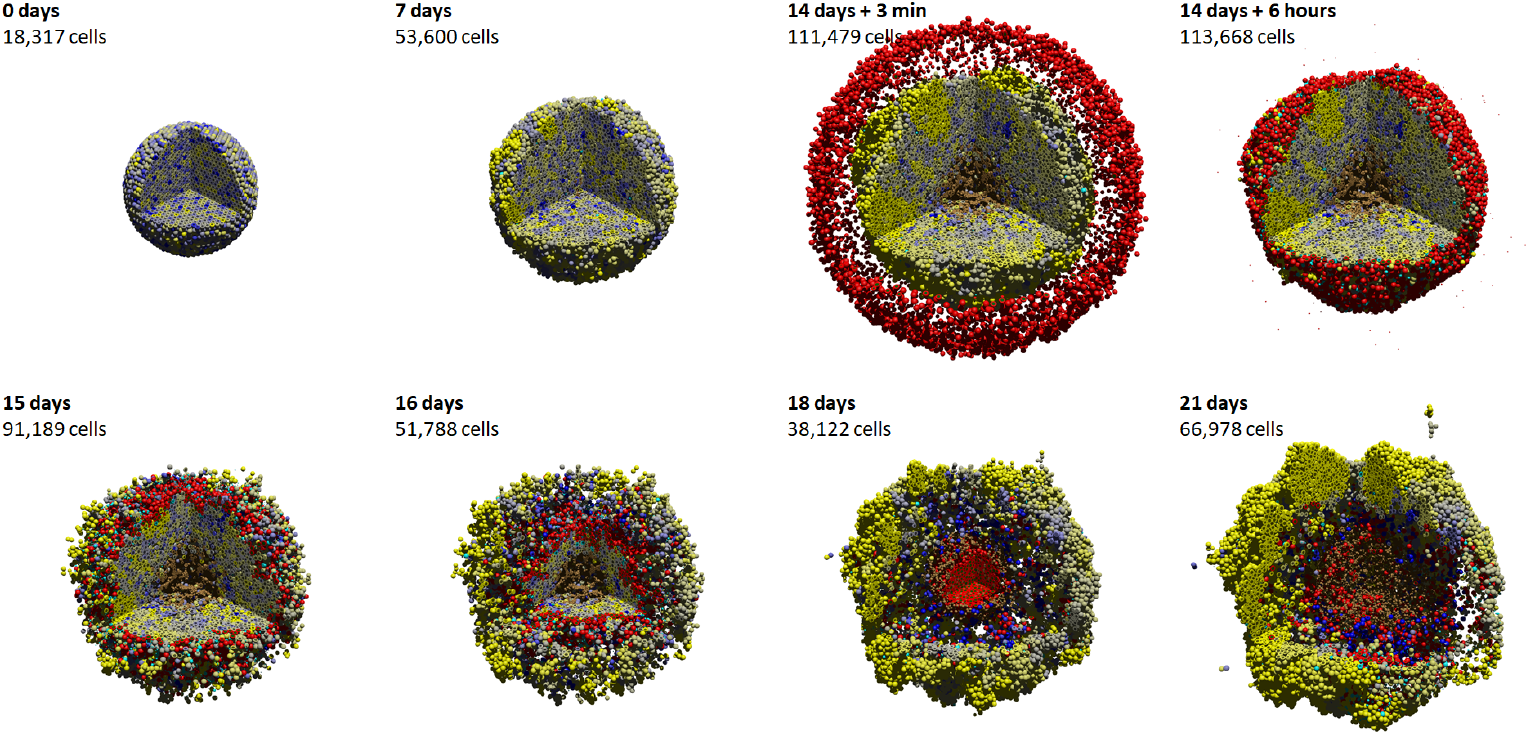
Cancer immunology example. In this 3-D example, each tumor cell secretes an immunostimulatory factor, and its immunogenicity is modeled as proportional to its mutant oncoprotein expression. (See the previous example in Fig. 9.) After 14 days, red immune cells perform a biased random walk towards the immunostimulatory factor, test for contact with cells, form adhesions, and attempt to induce apoptosis for cells with greater immunogenicity. The immune cells successfully attack the tumor initially, leading to partial regression; apoptotic cells are cyan. But strong homing towards gradients of the immunostimulatory factor causes immune cells to “pass” some cells at the outer edge, leading to tumor regrowth. Eventually, immune cells leave the necrotic regions and press their attack on the tumor. This highlights the importance of stochasticity in immune cell movement in mixing with the tumor cells for a more successful immune response. A video is available at S8 Video.

In [20,21], we used high-throughput computing to further expand this investigation to explore the impact of stochastic migration and tumor-immune cell adhesion dynamics on the treatment efficacy.

## Availability and Future Directions

### Getting started as a new PhysiCell user/developer

#### Necessary tools

Users need a working `g++` development environment with support for OpenMP and Makefiles (or a reasonably compatible 64-bit C++11 compliant compiler). (Future releases will likely include compile options using cmake.) We provide tutorials to set up a `g++` environment in Windows and OSX at:

##### OSX via Homebrew

http://www.mathcancer.org/blog/setting-up-gcc-openmp-on-osx-homebrew-edition/

##### OSX via MacPorts

http://www.mathcancer.org/blog/setting-up-gcc-openmp-on-osx-macports-edition/

##### Windows via mingw-w64

http://www.mathcancer.org/blog/setting-up-a-64-bit-gcc-environment-on-windows/

Mac OSX users should use the tutorials above; the version of `g++` included with XCode does not support OpenMP. Moreover, OSX users need to set a PHYSICELL_CPP environment variable as noted in the Homebrew and MacPorts tutorials.

Alternatively, users can code within a virtual machine using the PhysiCell virtual appliance. See the tutorial at http://www.mathcancer.org/blog/getting-started-with-a-physicell-virtual-appliar

#### Downloading PhysiCell

Users can download a PhysiCell release at:

##### SourceForge

https://sourceforge.net/projects/physicell/

The green “download” button will download the most recent source file (as a zip file). You can get the most recent virtual appliance by browsing to: https://sourceforge.net/projects/physicell/files/PhysiCell/ then browsing a recent release directory (e.g., PhysiCell 1.2.2), and downloading the ova file.

##### GitHub

https://github.com/MathCancer/PhysiCell/releases/latest Download either `PhysiCell_V.x.y.z.zip` (source) or `PhysiCell.x.y.z.ova` (virtual appliance).

PhysiCell is licensed under the (3-clause) BSD license, which is compatible with commercial products and can be included in GPL-licensed projects.

##### Learning to use PhysiCell

PhysiCell’s main project website can be found at http://PhysiCell.MathCancer.org.

Each PhysiCell download comes with a Quickstart guide (`Quickstart.pdf`) in the main directory (see S3 Text). We recommend running through this guide to populate, compile, run, and visualize your first projects. (The four examples in Additional PhysiCell examples are included as sample projects, with compile instructions in the Quickstart; see S3 Text.)

A full User Guide (see S2 Text) is included in the documentation directory of every PhysiCell download. See `documentation/User_Guide.pdf`. The User Guide includes full details on how to access and modify each cell’s `Phenotype`, how to write custom fuctions and data elements, and how to interact with the BioFVM microenvironment [6].

Users creating their own simulation projects should start with the 2-D and 3-D template projects. User-created code should be placed in the custom_modules directory; users should not modify functions in `core` (core PhysiCell functionality) or `modules` (PhysiCell-provided, non-core functions). See the Quickstart (S3 Text) and User Guide (S2 Text) for further details.

We post tutorials at our PhysiCell blog, available at http://mathcancer.org/blog/physicell-tutorials.

Check there for tips and tricks to building simulators, and visualizing results. We also frequently post updates on PhysiCell on Twitter, under the hashtag `#PhysiCell`.

### Getting started as a new PhysiCell contributor

We welcome new functionality and bug fix contributions from the community. No special tools or libraries are required for PhysiCell development aside from a C++11/OpenMP compliant compiler, `make`, and a copy of the PhysiCell source. Developers should fork the current development version of the PhysiCell repository (available on GitHub at https://github.com/MathCancer/PhysiCell), develop their contributions (either in the `modules` directory as new functions, named `as PhysiCell. [name].cpp`, or as fixes to existing files), and then submit a pull request. Users should also email the PhysiCell mailing list when submitting code contributions.

Project coding conventions can be found at http://www.mathcancer.org/blog/mathcancer-c-style-and-practices-guide/.

### Getting help

Users encountering problems in compiling PhysiCell projects should first consult the FAQ page at: http://www.mathcancer.org/blog/common-problems-and-solutions/

Users (and developers) can join the PhysiCell mailing list to ask the community questions or discuss ideas, at. Users can submit bug reports at the GitHub issue tracker: https://github.com/MathCancer/PhysiCell/issues

### Limitations and potential solutions

#### Scientific limitations

BioFVM does not implement advective transport as of Version 1.1.7, and so PhysiCell cannot readily be applied to advection-dominated problems without user-supplied advection solvers. We also have not yet written a model of extracellular matrix; users could add this effect by introducing custom data that (1) “anchors” each cell to a position in space beyond cell-cell mechanical operations, and (2) slowly evolve these “anchor points” to model ECM rearrangement. We are currently exploring this approach to model liver parenchyma, and we will post sample codes as the work progresses. Alternatively, users could introduce an independent finite element mesh for the ECM, evolve it under (e.g., viscoelastic) laws, and attach cells to the nearest lattice site. As a simpler approach, users could include a non-diffusing ECM substrate (in BioFVM), which could be used to reduce the cells’ velocities (as extra drag). Vary the cell-cell mechanics parameters in `phenotype.mechanics` with the ECM density for this approach.

PhysiCell’s cell-centered approach may not be ideal for application to some morphogenesis problems; see the recent work by Osborne and co-workers that compared several discrete modeling frameworks in morphogenesis test problems [47]. PhysiCell does not model cell morphology, and so it cannot directly model cell elongation processes needed in some morphogenesis problems. However, users could potentially model elongated cells as two or more agents, and introduce manual adhesive links similar to our “biorobots” examples. PhysiCell has not implemented cell fusion (another mechanism of morphogenesis), but users could manually implement this by (1) testing for cell-cell contact, (2) moving one cell to the center of volume of the two cells, (3) updating that cell’s volume to include both original cells’ volumes, and (4) deleting the second cell. PhysiCell has not yet written functions to deal with polarized cell-cell adhesion, or to update its orientation. Thus, PhysiCell may need further development and user contributions to model the full spectrum of morphogenetic mechanisms.

We note that PhysiCell can manually implement molecular-scale biology as ODEs (e.g., as in the drug damage example) via custom functions and data, but this is currently difficult to scale to large systems of ODEs with many parameters. Moreover, while we have provided examples of user-defined cell cycle models and other custom functions in the User Guide, sample codes, and tutorials, we do not provide templates for such functions beyond the documentation. We are currently testing methods to support SBML specification of ODE models in the cell agents (e.g., via libRoadrunner [48]), but until SBML is supported, we do not provide templates to integrate ODE models into PhysiCell functions to alter cell phenotype.

PhysiCell does not currently provide direct methods to model contact-based cell-cell interactions; these can be manually implemented as we did in the immune and biorobots examples. We also do not yet provide alternative mechanics models (e.g., viscoelastic) out-of-the-box, but users can design their own cell velocity functions as needed to replace the default mechanics. We have not yet implemented direct calculations of cell pressures and strains, so this could hinder some applications in mechanobiology. Users could implement this by creating a custom velocity update function that calculates pressures and strains from the potential functions. A preliminary version—calculated from the “resistance” potential functions similarly to [49]—is included as `state.basic_pressure`. See the User Guide (S2 Text) for more details.

#### Software limitations

PhysiCell is intended to function as a modular engine and to interact with standardized data (e.g., MultiCellDS [22]), but it has not yet fully implemented MultiCellDS import and export. Graphical simulation design tools, data visualization tools, and analysis tools are needed to widen its accessibility beyond seasoned C++ programmers and reduce its learning curve. In the future, PhysiCell should switch from Makefiles to CMake to facilitate simpler cross-platform compiling.

### Future improvements

#### Numerical improvements

The biggest performance bottleneck is cell-cell interaction testing: cell volume can vary by a factor of 100, and hence the cell diameter (and interaction distance) can vary by a factor of 50. The number of cells in the list of interacting neighbors 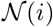 scales inversely with the minimum cell volume; see Expanded computational cost estimates. Future versions of PhysiCell will introduce a nested mesh interaction testing structure to more accurately estimate 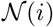 in regions with small cells. Although PhysiCell’s design objectives are not focused on larger simulations with 10^8^-10^9^ cells, extension to supercomputers by wrapping the code in MPI with appropriate data mirroring could be useful. Likewise, the algorithms used in BioFVM could be implemented on a graphics card via OpenCL or OpenACC. We will explore these approaches in the future.

#### Scientific improvements

We will develop SBML (systems biology markup language) importers for molecular-scale biology, most likely by the C code generation features in COPASI [50] or libRoadrunner [48]. This will make it straightforward to integrate systems of ODEs to individual cells. We will add new cycle models for flow cytometry-driven problems (G1, S, and G2/M phases). We plan to add more advanced cell mechanics models (e.g., as in [9, 51,52]), and to extend PhysiCell’s included standard functions to include extracellular matrix mechanics. We also plan to introduce built-in functions for polarized cell adhesion and updating the cells’ orientations. As our PhysiCell-based projects progress, we will “upstream” our new functions to PhysiCell as new optional modules. We are currently testing and refining the elastic-based cell adhesions in Additional PhysiCell examples for inclusion as standard models, and we are also developing vascularization libraries as part of our work to simulate metastatic breast cancers injected in mammary fat pads.

#### User-focused improvements

In the coming months, we will continue publishing blog posts and code samples at http://MathCancer.org/blog/physicell-tutorials/. We will create pre-compiled clients that can initiate simulations based upon a digital snapshot (intitial arrangement of cells) and digital cell lines (self-contained, model-independent sets of cell phenotype data), using the emerging MultiCellDS standard [22, 23]. We will develop support to read parameters from XML configuration files, and we plan to offer PhysiCell as a compiled, linkable library. Moreover, we plan to develop user-friendly, web-based applications of PhysiCell to simulate the multicellular response to diffusing engineered nanoparticles [53].

## Supporting Information

### S1 Text

#### Supplemental information

Extensive supplemental information including: full mathematical model details, supporting literature, and reference parameter values for breast epithelial cells; expanded numerical implementation details; convergence and validation testing results; full parameter values for the main tests; and an expanded feature comparison of PhysiCell and other 3-D multicellular simulation platforms.

### S2 Text

#### User Guide

User Guide for PhysiCell 1.2.2. Includes full API documentation and examples.

### D3 Text

#### Quickstart guide to PhysiCell

A fast guide to building, running, and visualizing your first PhysiCell simulations. We recommend that new users start with this guide.

### S1 Video

#### Deterministic 3-D hanging drop spheroid simulation

3-D simulation of 18 days of hanging drop tumor spheroid growth from 2300 cells to 1.2 million cells, using the deterministic necrosis model. Available at: https://www.youtube.com/watch?v=WMhYW9D4SqM and https://doi.org/10.6084/m9.figshare.5716600

### S2 Video

#### Stochastic 3-D hanging drop spheroid simulation

3-D simulation of 18 days of hanging drop tumor spheroid growth from 2300 cells to 1 million cells, using the stochastic necrosis model. Available at: https://www.youtube.com/watch?v=xrOqqJ_Exd4 and https://doi.org/10.6084/m9.figshare.5716597

### S3 Video

#### Deterministic 3-D ductal carcinoma in situ (DCIS) simulation

3-D simulation video of 30 days of DCIS growth in a 1 mm length of breast duct, using the deterministic necrosis model. Available at: https://www.youtube.com/watch?v=ntVKOr9poro and https://doi.org/10.6084/m9.figshare.5716480

### S4 Video

#### Stochastic 3-D ductal carcinoma in situ (DCIS) simulation

3-D simulation video of 30 days of DCIS growth in a 1 mm length of breast duct, using the stochastic necrosis model. Available at: https://www.youtube.com/watch?v=-lRot-dfwJk and http://dx.doi.org/10.6084/m9.figshare.5721088.v1

### S5 Video

#### 2-D biorobots simulation

2-D simulation of the “biorobots” example, showing a synthetic multicellular cargo delivery system. Available at: https://www.youtube.com/watch?v=NdjvXI_x8uE and https://doi.org/10.6084/m9.figshare.5721136

### S6 Video

#### 2-D biorobots, applied to cancer therapeutics delivery

2-D simulations of the “biorobots” adapted for use as a cancer treatment, where cargo cells detach and secrete a therapeutic once reaching hypoxic tissues. Available at: https://www.youtube.com/watch?v=wuDZ40jWM and https://doi.org/10.6084/m9.figshare.5721145

### S7 Video

#### 2-D simulation of a heterogeneous tumor

2-D simulation of a tumor whose heterogeneous oncoprotein expression drives proliferation and selection. Available at: https://www.youtube.com/watch?v=bPDw6l4zkF0 and https://doi.org/10.6084/m9.figshare.5721151

### S8 Video

#### 3-D simulation of a tumor immune response

3-D simulation of immune cells attacking a tumor with heterogeneous proliferation and immunogenicity. Available at: https://www.youtube.com/watch?v=nJ2urSm4ilU and https://doi.org/10.6084/m9.figshare.5717887

## Acknowledgments

We thank the Breast Cancer Research Foundation and the Jayne Koskinas and Ted Giovanis Foundation for Health and Policy for generous support. Partial support was provided by the USC James H. Zumberge Research and Innovation Fund (2012 Large Interdisciplinary Award), the National Institutes of Health (5U54CA143907,1R01CA180149), and the National Science Foundation (1720625). We thank David B. Agus and the USC Center for Applied Molecular Medicine for generous institutional support. We thank Margy Villa for her assistance in early testing, and Edwin Juarez and Rishi Rawat for useful discussions. We greatly appreciate the deep discussions with the MultiCellDS review panel in developing standardized cell cycle and death models. We thank Jakob Kather and Jan Poleszczuk for discussions on cancer immunology. We thank Nathan Choi for his 3-D hanging drop spheroid work in Figure 2.

